# Potent SARS-CoV-2 neutralizing antibodies selected from a human antibody library constructed decades ago

**DOI:** 10.1101/2020.11.06.370676

**Authors:** Min Qiang, Peixiang Ma, Yu Li, Hejun Liu, Adam Harding, Chenyu Min, Lili Liu, Meng Yuan, Qun Ji, Pingdong Tao, Xiaojie Shi, Zhean Li, Fulian Wang, Yu Zhang, Nicholas C. Wu, Chang-Chun D. Lee, Xueyong Zhu, Javier Gilbert-Jaramillo, Abhishek Saxena, Xingxu Huang, Hou Wang, William James, Raymond A. Dwek, Ian A. Wilson, Guang Yang, Richard A. Lerner

## Abstract

Combinatorial antibody libraries not only effectively reduce antibody discovery to a numbers game, but enable documentation of the history of antibody responses in an individual. The SARS-CoV-2 pandemic has prompted a wider application of this technology to meet the public health challenge of pandemic threats in the modern era. Herein, we used a combinatorial human antibody library constructed 20 years before the COVID-19 pandemic to discover three highly potent antibodies that selectively bind SARS-CoV-2 spike protein and neutralize authentic SARS-CoV-2 virus. Compared to neutralizing antibodies from COVID-19 patients with generally low somatic hypermutation (SHM), these antibodies contain over 13-22 SHMs, many of which are involved in specific interactions in crystal structures with SARS-CoV-2 spike RBD. The identification of these somatically mutated antibodies in a pre-pandemic library raises intriguing questions about the origin and evolution of human immune responses to SARS-CoV-2.

## INTRODUCTION

The global spread of SARS-CoV-2, a novel coronavirus and cause of the coronavirus disease 2019 (COVID-19), poses an unprecedented health crisis and was declared a pandemic by the World Health Organization on March 11, 2020 (*1*). To date, over 40 million individuals have been infected with over 1 million deaths (https://covid19.who.int/) with no vaccines or specific antiviral drugs yet approved. Monoclonal antibodies (mAbs) targeting the viral spike glycoprotein (S) have been shown to have excellent neutralization efficacy in previous treatment of SARS, MERS and Ebola infections, and therefore are of particular interest to combat the current pandemic (*2, 3*). Since the COVID-19 outbreak, the spike glycoprotein has been the main target for development of therapeutic mAbs (*4, 5*). Most neutralizing antibodies (NAbs) bind to the receptor binding domain (RBD) of the S protein (*6*), although some also bind to the N-terminal domain (NTD) (*7, 8*). NAbs have been derived from multiple sources, including memory B cells from SARS-CoV-2 convalescent patients (*3, 9, 10*), previous SARS patients (*11*), immunized humanized H2L2 mice (*12*), alpaca nanobodies (*13*), single domain human antibodies from a pre-established library (*14*), and phage display antibody libraries (*4, 15, 16*).

Antibody generation is an evolutionary process of mutation and selection from the B cell repertoire. The combinatorial antibody library technology allows the same evolutionary process to be performed *in vitro* as it restores the "fossil record" of an individual's antibody response in a test tube (*17*). Random coupling of V_H_ and V_L_ sequences in scFv libraries greatly expands diversity, thereby allowing for selection of novel antibodies with high binding affinity and neutralization efficacy (*18–22*).

Here, we report the selection and characterization of three potent SARS-CoV-2 antibodies, S-E6, S-B8 and S-D4, from a pre-pandemic human combinatorial antibody library established in 1999 that target the spike RBD and compete with *h*ACE2 receptor (*23*). This study provides further evidence that a combinatorial antibody library with an unprecedented diversity can mimic the selection process of natural immunity, permit detection of rare spike-targeting antibodies with higher somatic hypermutation, and allow for selection of binding molecules with chemistries beyond those accessible during *in vivo* selection.

## RESULTS

### Selection of antibodies against SARS-CoV-2 spike RBD

We constructed and overexpressed the SARS-CoV-2 spike RBD (S-RBD) linked to human Fc (*h*Fc) with a thrombin digestion site. After affinity purification, recombinant SARS-CoV-2 S-RBD was biotinylated, immobilized on streptavidin-coated magnetic beads, and panned against a combinatorial scFv antibody phage library containing 10^11^ members generated from peripheral blood mononuclear cells (PBMC) of 50 healthy donors in 1999 (*23, 24*). In the first two rounds, a pH 2.2 glycine-HCl solution was used to elute antibody-displaying phagemids bound to S-RBD. To enrich for antibodies that compete with *h*ACE2, a “function-guided enrichment” strategy was used in the third round, where recombinant *h*ACE2-ECD protein was used to elute S-RBD binding phagemids. After three rounds of panning, S-RBD-specific antibodies were enriched (Figure 1A) and 22 unique antibodies were selected that specifically bound to S-RBD-*h*Fc (Figure 1B) (*24*).

**Figure 1.**
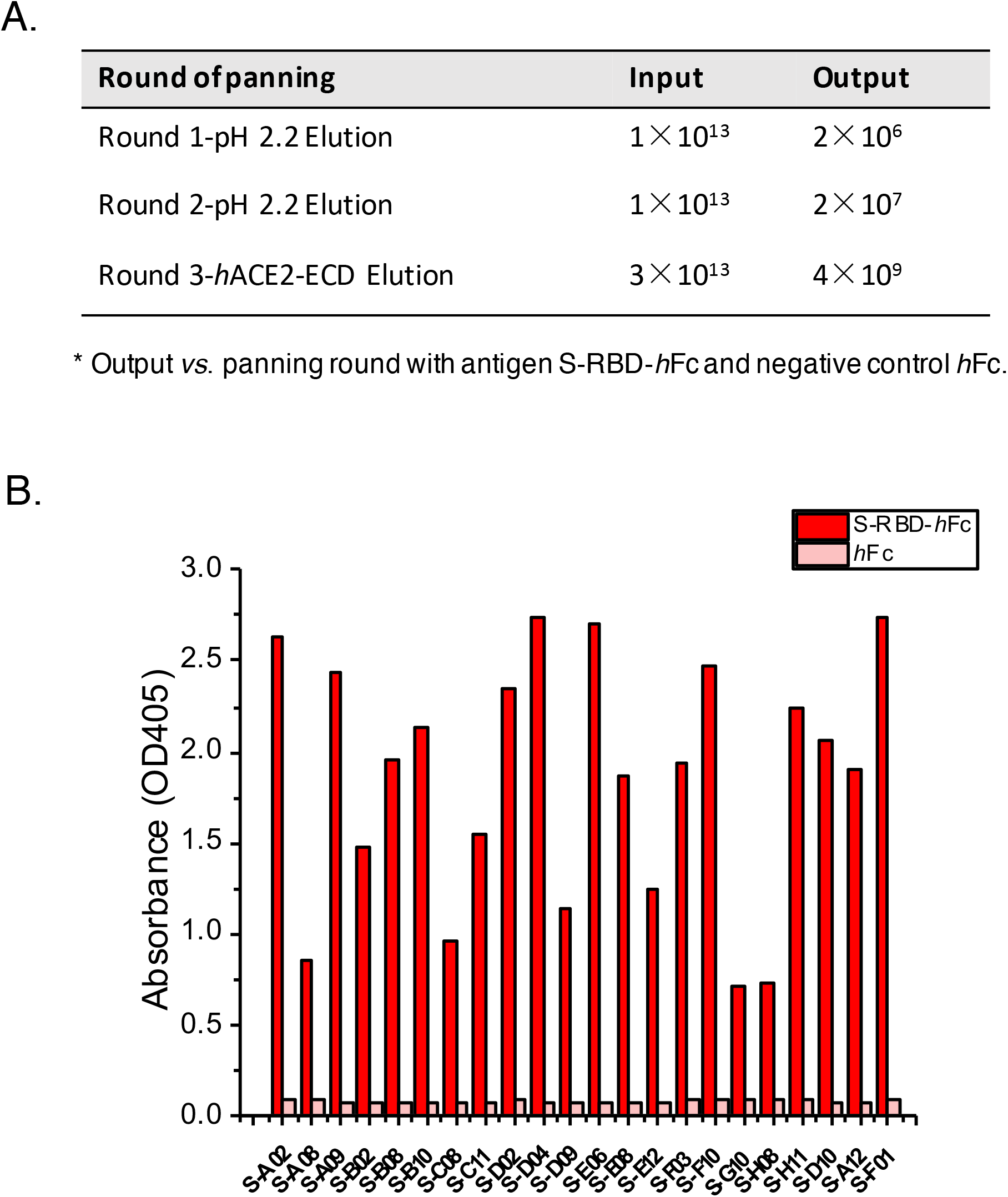
Selection of scFv antibodies targeting spike protein. (**A**) Output vs. panning round for the antigen S-RBD-*h*Fc during three rounds of screening. (**B**) Phage ELISA results of 22 unique antibodies with positive readouts (OD405 ratio S-RBD-*h*Fc/*h*Fc > 2).

Next-generation sequencing of the library revealed that 92% of human heavy-chain IGHV and 89% of the light-chain (IGLV and IGKV) germlines were covered, when aligned to the IMGT (international ImMunoGeneTics) database (Supplementary Figure 1), enabling screening of antibodies encoded by diverse germlines.

### Selected anti-S-RBD antibodies retain binding to full-length spike

The scFv antibodies were then converted to full-length monoclonal antibodies (mAbs) by cloning into a human IgG4e1(S228P) vector. HEK293F cells were adapted for expression of combinatorial antibodies that were secreted into culture supernatants (*24*). Three of the best performing antibodies, S-B8, S-D4 and S-E6, were purified to homogeneity with yields of 8.1, 9.6 and 17 mg/L, respectively, whereas S-RBD-*h*Fc (IgG1) was 58 mg/L (Supplementary Figures 2 & 3). To characterize interactions between the anti-S-RBD antibodies and full-length spike, HEK293T cells were transiently transfected with either SARS-CoV-2 spike-P2A-EGFP or SARS spike-P2A-EGFP (*24*). Flow cytometry (FACS) showed that all three antibodies in full-length-IgG4 format retained their ability to bind full-length SARS-CoV-2 spike (Figure 2A & 2D) with no cross-reactivity with SARS-CoV spike (Figure 2B) or non-transfected cells (Figure 2C).

**Figure 2.**
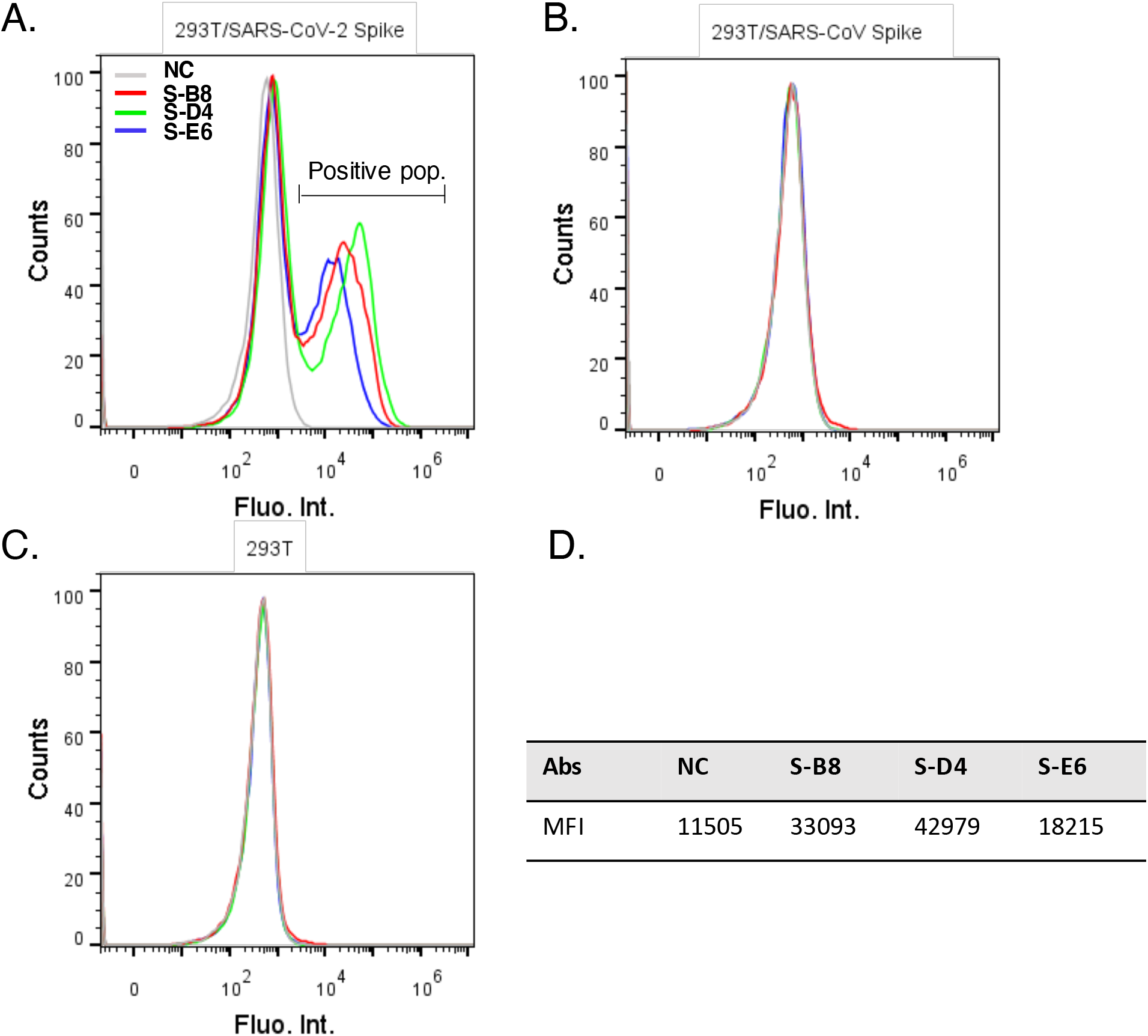
Analysis of antibody binding to cell surface-expressed trimeric spike protein. (**A**) HEK293T cells transfected with expression plasmid encoding the full-length spike of SARS-CoV-2 were incubated with purified IgG4 antibody and stained with PE labeled anti-human IgG4 Fc secondary antibody, then analyzed by FACS. Positive binding cells populations were labeled as positive pop. (**B**) FACS of antibodies binding to SARS-CoV spike. (**C**) FACS of antibodies binding to non-transfected HEK293T cells. Cells stained with only secondary antibody were used as negative control (NC). (**D**) Mean fluorescent intensity (MFI) of antibodies for SARS-CoV-2 spike binding, *i.e*. positive population area in **A**.

### Antibody binding and competition with *h*ACE2-ECD to SARS-CoV-2 S-RBD

To assess neutralization potential of the mAbs, we investigated their ability to compete with ACE2-ECD for S-RBD binding by ELISA (*15, 25*). S-B8, S-D4 and S-E6 all competed strongly with *h*ACE2-ECD in a dose-dependent manner, with IC_50_ values of 12.9±1.5 nM, 7.1±0.4 nM, and 12.2±0.7 nM, respectively (Figure 3A). Kinetic parameters of on-rate (*k*_on_), off-rate (*k*_off_), and dissociation constant (K_D_) for the three antibodies were then determined by biolayer interferometry (Figure 3B – 3E). S-B8, S-D4, and S-E6 showed K_D_ values of 170 pM, 10 pM and 110 pM, respectively, with S-D4 displaying the highest affinity due to its greatly reduced off-rate (Figure 3C).

**Figure 3.**
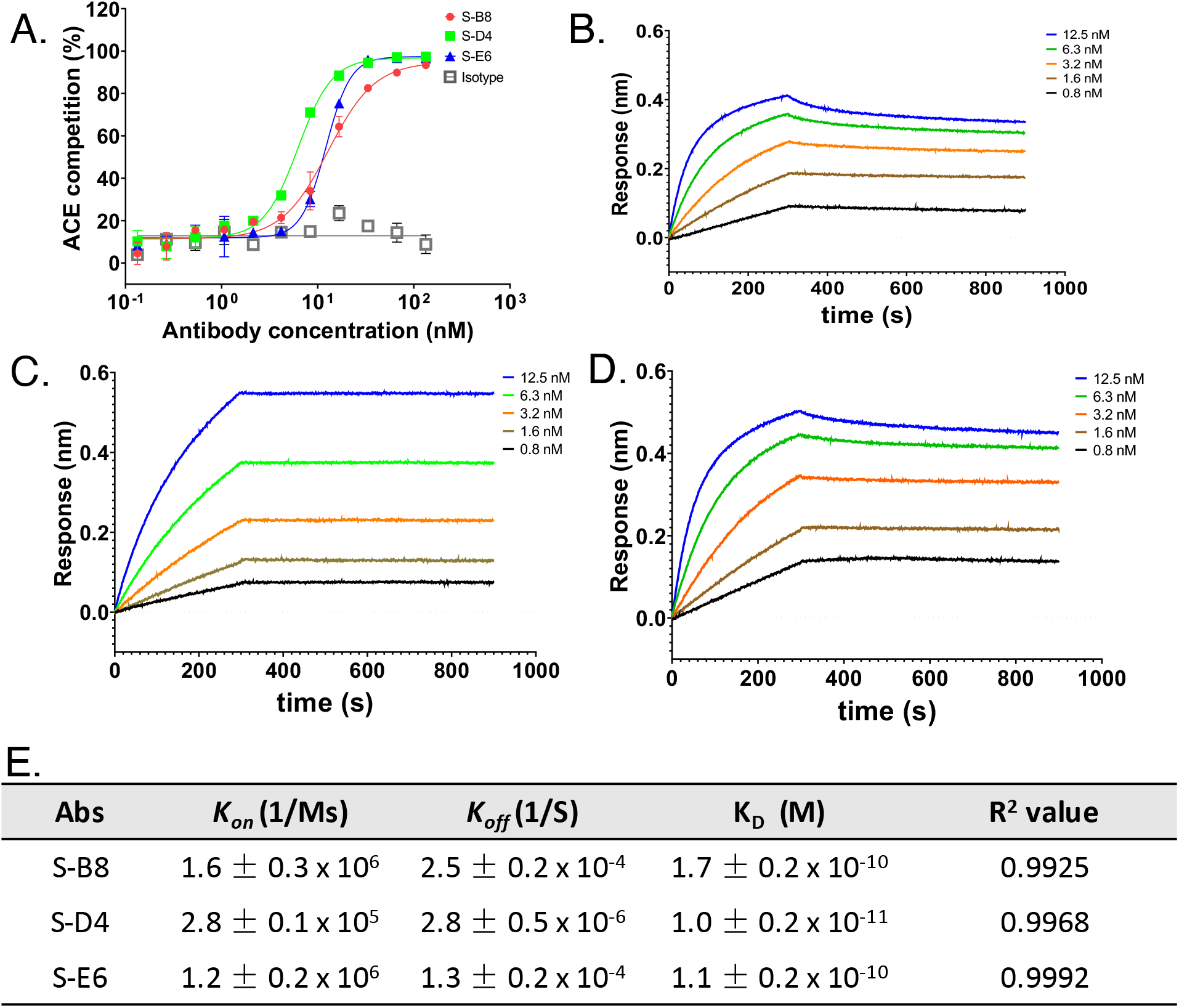
Competitive ELISA of antibodies with *h*ACE2 and binding kinetics to the spike protein. (**A**) The three antibodies were titrated for competition with *h*ACE2-ECD for binding to S-RBD, the fitting curves are shown. (**B-D**) Binding kinetics were measured by biolayer interferometry (BLI). Biotinylated S-RBD was loaded to the SA biosensor for detection of binding kinetics with S-B8 and S-E6, while S-RBD amine coupled to AR2G sensor was utilized for S-D4, with detection on Octet. All curves were fitted by a 1:1 binding model using the Data Analysis software (Forte Bio). (**E**) The association-rate (kon), dissociation-rate (koff) and dissociation constant (KD) of the three competitive antibodies are shown.

We also tested with natural mutants of SARS-CoV-2 spike proteins that have been clinically associated with more severe illness and longer hospital stays. Three mutants, *i.e.* D215H (mut 1), S247R (mut 2) and D614G (mut 3), isolated from patients requiring treatment in an intensive care unit (ICU) (*26*), all displayed similar binding by FACS to S-B8, S-D4, and S-E6 (Supplementary Figure 4), indicating their therapeutic potential against natural SARS-CoV-2 spike mutants in severely affected patients.

### Inhibition of cell-cell fusion induced by SARS-CoV-2 spike and *h*ACE2

The S2 subunit of the SARS-CoV-2 spike mediates membrane fusion in *h*ACE2 expressing cells and is essential for virus infection. *h*ACE2 binding to SARS-CoV-2 is stronger than to the SARS-CoV spike (K_D_ of 4.7 nM and 32 nM, respectively) (*27*). To test whether these antibodies could inhibit spike-mediated membrane fusion of cells, we established a cell-cell fusion assay using Vero cells overexpressing *h*ACE2 as target cells, SARS-CoV-2 spike-P2A-EGFP transient transfected HEK293F cells as effector cells, and SARS-CoV spike-P2A-EGFP cells as a negative control (*24*). Spike-expressing HEK293F cells were mixed with S-B8, S-D4 or S-E6 at 10 nM or 1 nM just before adding to the Vero cells and syncytium formation observed 6 hours later. The SARS-CoV-2 spike induced significant cell-cell fusion as manifested by formation of larger EGFP positive cells, whereas the SARS-CoV spike barely induced syncytium formation (Figure 4A). All three antibodies potently inhibited cell-cell fusion by SARS-CoV-2 at both 10 and 1 nM (Figure 4B, 4C & 4F). At 10 nM, S-D4 and S-E6 showed over 80% inhibition of cell-cell fusion, which was slightly greater than recombinant S-RBD; S-D4 and S-E6 were more potent than S-B8 at 1 and 10 nM (Figure 4D & 4E).

**Figure 4.**
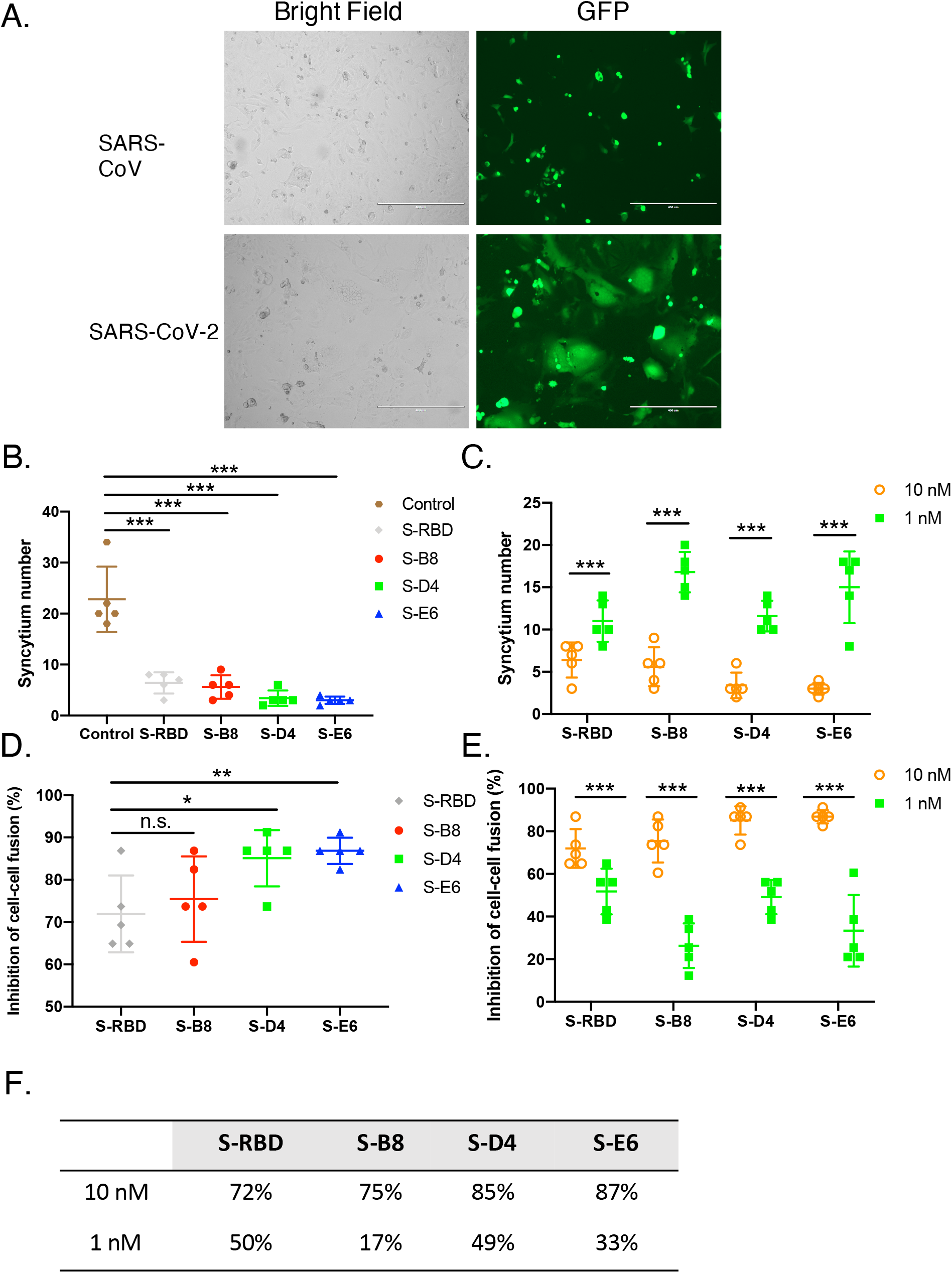
Inhibition of syncytium formation by the antibodies. (**A**) Representative images of SARS-CoV-2 and SARS-CoV spike-mediated syncytium formation with *h*ACE2 expressing cells 48 hours after co-culture. (**B**, **D**) Syncytium number calculation and inhibition rates when treated with 10 nM of *h*ACE2 competitive antibodies are shown. S-RBD was used as the positive control. (**C, E**) Syncytium number and inhibition rates treated by antibodies and S-RBD at different concentrations are shown. The inhibition rates at 10 nM and 1 nM are summarized in **F**. Bars=400 μm. Error bars indicate SD, \**P* < 0.05, \*\**P* < 0.01, \*\*\**P* < 0.001, determined by Student’s T-test.

### Inhibition of SARS-CoV-2 pseudovirus and authentic virus

To test neutralization against SARS-CoV-2 virus, we assessed the antibodies in a pseudovirus (PSV) infection assay. Pseudotyped particles were pre-incubated with S-B8 and S-D4 (from 200 nM to 200 fM) and S-E6 (200 nM to 6.3 fM), followed by infection of HEK293T/*h*ACE2 cells (*24*). Luciferase activity resulting from infection was determined at 60 h post transfection. All three antibodies showed potent neutralization against PSV infection in a dose-dependent manner that went to completion. The NT_50_ values of S-B8, S-D4, and S-E6 in pseudovirus neutralization were determined to be 2.2±0.2 nM, 0.48±0.03 nM, and 0.025±0.002 nM, respectively (Figure 5A) in a 1:1 interaction model with HillSlopes near 1.0 (Figure 5B). We next tested antibody neutralization of authentic SARS-CoV-2 virus [BetaCoV/Australia/VIC01/2020; GenBank MT007544.1 (Victoria/01/2020)] (*24*). Twenty hours after infection, intracellular virus was visualized and quantitated as percent infectivity. S-D4 and S-E6 were capable of fully blocking infection by authentic virus, while S-B8 was less potent (Figure 5C) with NT_50_ values for S-B8, S-D4 and S-E6 of 16 nM, 0.7±0.08 nM and 0.25±0.05 nM, respectively (Figure 5D).

**Figure 5.**
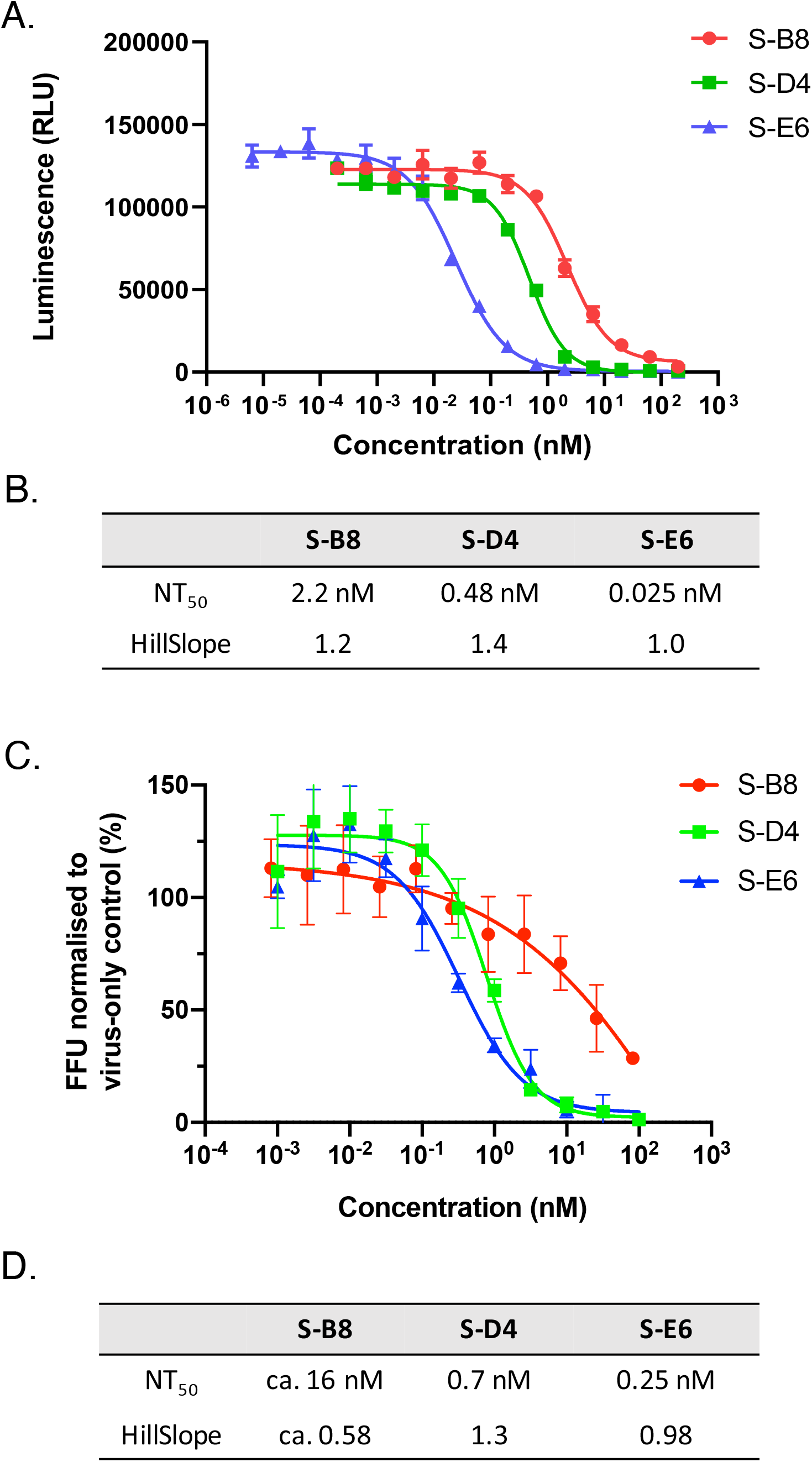
Neutralization assay for the *h*ACE2 competitive antibodies. **(A)** Neutralization ability of the three *h*ACE2 competitive antibodies to SARS-CoV-2 pseudovirus was tested and fitted. (**C)**A microneutralization assay was adopted for testing of the three antibodies. NT50 and HillSlope for each antibody are summarized in **B** and **C**.

### S-B8 and S-E6 bind the RBD and sterically block ACE2 binding

To elucidate the molecular recognition of S-B8 and S-E6 for SARS-CoV-2 S-RBD, x-ray structures of Fab/RBD complexes were determined to 2.25 and 2.70 Å, respectively (Supplementary Table 1). Fab S-B8 and S-E6 bind the receptor binding site (RBS) with different approach angles (Figure 6A) and sterically compete with ACE2 for RBD binding, consistent with the competition assay (Figure 3A). S-B8 interacts mainly using its heavy chain, which contributes 73% of the buried surface area (BSA, 737 of 1010 Å^2^) (Figure 6B) and 12 of 16 polar contacts (Supplementary Table 2). S-E6 predominately uses its light chain, which contributes 63% of the BSA (530 of 847 Å^2^) and 16 of 19 polar contacts (Supplementary Table 2). Light chain dominant interactions are less common in antibodies (*28, 29*).

**Figure 6.**
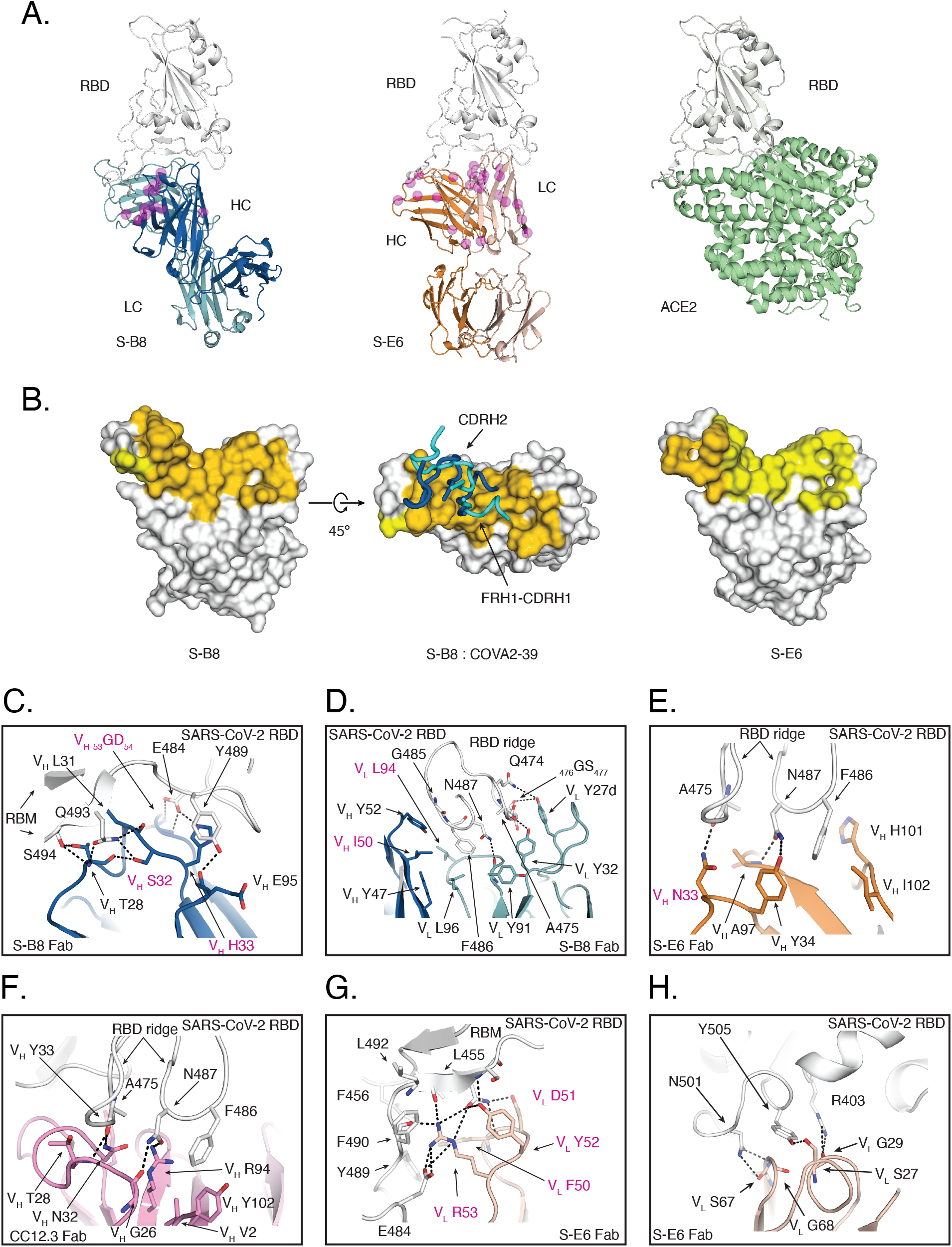
Structural characterization of S-B8 and S-E6 integrating with SARS-CoV-2 S-RBD. Crystal structures are shown in ribbon representation with residues of interest in stick mode. The epitope surface on the RBD involved in interaction with the heavy and light chains of the antibodies are in orange and yellow, respectively. S-RBD is shown in white, S-B8 in blue and light blue for heavy and light chains, and S-E6 heavy and light chains in orange and pink, *h*ACE2 is shown in green. SHM residues are shown as semi-transparent magenta spheres and highlighted with magenta labels in **C-H**. (**A)** Structure comparison of S-B8 and S-E6 compared to *h*ACE2 binding to the RBD in the same relative view. (**B)** Surface representation of S-RBD epitope residue interactions with S-B8 and S-E6. FRH1-CDRH1 and CDRH2 from both S-B8 (blue) and COVA2-39 (cyan, PDB 7JMP) are shown for comparison. (**C**) S-B8 CDRH1 and CDRH2 interaction with RBD. (**D)** Interaction between S-B8 and RBD ridge. (**E)**Interaction between S-E6 and RBD ridge. (**F**) Comparison to IGHV3-53 binding mode A. CC12.3 (pink for heavy chain and light pink for light chain) in complex with SARS-CoV-2 S-RBD (PDB 6XC7) illustrating the hydrogen bonding between the 32NY33 motif and S-RBD. (**G**) Interaction between S-E6 and RBM mid-region. (**H**) Interaction between S-E6 and RBM on the opposite side of the S-RBD ridge.

IgBLAST analysis (*30*) suggests S-B8 is derived from IGHV3-66, a germline that is highly similar to IGHV3-53 (Supplementary Figure 5). We previously reported that IGHV3-53 antibodies from convalescent patients, with minimal SHM and high potency, have two key germline motifs in CDRH1 and CDRH2 that are primarily used for recognition of SARS-CoV-2 S-RBD, namely _32_NY_33_ and _53_SGGS_56_ (*31, 32*). In addition, two very distinct binding modes (A and B) are observed for IGHV3-53/3-66 antibodies depending on CDRH3 length (*31–33*). However, in S-B8, _32_NY_33_ in CDRH1 is mutated to _32_SH_33_ and _53_SCGS_56_ (_53_TGGT_56_ in COVA2-39) in CDRH2 to _53_GDGN_56_ (Supplementary Figures 6 & 7). Intriguingly, CDRH1 and CDRH2, as well as FRH1, of S-B8 still bind to a similar region on SARS-CoV-2 S-RBD to that of binding mode B (Figure 6B) (*31*). The _32_SH_33_ in S-B8 is part of a type I beta-turn (Figure 6C). V_H_ S32 interacts with RBD Q493 and its side-chain hydroxyl hydrogen bonds to the V_H_ T28 carbonyl oxygen. The V_H_ H33 imidazole forms a salt bridge with RBD E484 and a π-π interaction with Y489. The V_H 53_GD_54_ backbone in CDRH2 also forms two hydrogen bonds with E484, and V_H_ T28 and L31 make four hydrogen bonds with Q493 and S494 (Figure 6C, Supplementary Table 2).

F486 in the S-RBD ridge region is buried in a hydrophobic pocket (V_H_ W47, V_H_ I50, V_L_ Y91, V_L_ L94, V_L_ L96) between the heavy and light chains, while _485_GF_486_ and _476_GS_477_ on the RBD ridge interact with V_H_ Y52 and V_L_ Y27d *via* π-π interactions (Figure 6D). Of note, F486 is also buried in a pocket at the heavy-light chain interface in COVA2-39, which is an IGHV3-53 antibody, as well as other antibodies that bind in RBS-B mode (*34*). Altogether, 19 of 29 S-B8 epitope residues are shared with 19 of 21 COVA2-39 epitope residues, with16 corresponding to ACE2 binding residues (Supplementary Figure 8).

### S-E6 interaction with SARS-CoV-2 S-RBD

S-E6 is an IGHV4-31 antibody. Interestingly, SHM introduces a 33NY34 sequence in a similar position to the _32_NY_33_ motif in CDRH1 in IGHV3-53/3-66 antibodies (Supplementary Figure 6) that interact with the same RBD site but in a different orientation compared to _32_NY_33_ of IGHV3-53 binding mode A (*34*). Nevertheless, V_H_ N33 still hydrogen bonds with RBD A475 carbonyl (Figure 6E), as does V_H_ N32 of IGHV3-53 in binding mode A (Figure 6F). V_H_ Y34 and V_H_ A97 form two hydrogen bonds with N487 of the S-RBD (Figure 6E), which differ from Y33 in IGHV3-53 antibodies (Figure 6F). F486, along with N487, interact with a hydrophobic pocket formed by V_H_ Y34, A97, H101 and I102 of S-E6 and also make π-π and cation-π interactions (Figure 6E). However, the S-E6 light chain contributes the majority of the buried surface with RBD. CDRL2 _50_FDYR_53_ interact with the receptor binding motif (RBM) *via* multiple polar interactions (8 hydrogen bonds and 3 salt bridges) to E484, F490, L492, Q493 and S494 (Figure 6G, Supplementary Table 2). Moreover, V_L_ F50 interacts with a nearby hydrophobic patch formed by L455, F456 and Y489 (Figure 6G), and V_L_ S27, G29, S67 and G68 form five hydrogen bonds with R403, N501 and Y505 on the other side of the RBS ridge (Figure 6H, Supplementary Table 2).

### SHM residues form specific interactions with the RBD

Unlike most RBD-targeting neutralizing antibodies isolated from COVID-19 patients which have minimal SHM, the antibodies derived from this combinatorial antibody library are more highly mutated; S-B8 and S-E6 contain 13 and 22 SHM residues, respectively, several of which are in the antibody paratope (Figure 6 and Supplementary Figure 6), including V_H 31_LSH_33_, V_H 50_IT_51_, V_H 53_GD_54_, V_H_ N_56_, V_H_ D_58_ and V_L_ L94 in S-B8, and V_H_ N_33_, V_L_ V_39_, V_L 50_FDYR_53_ and _65_TR_66_ in S-E6 (Figure 6, C – G, and Supplementary Figure 6). In summary, several SHM residues appear to be critical for interaction with SARS-CoV-2 RBD, despite the antibody libraries being generated 20 years ago, implying the possibility that the eliciting antigen was structurally very similar to the SARS-CoV-2 RBD or there is rare but fortuitous cross-reactivity with another antigen.

### Antibody autoreactivity

To investigate the origin of the three antibodies, a HEp-2 autoreactivity assay was performed. Neither S-D4 nor S-E6 showed a positive signal in the assay, suggesting that they are not derived from auto-immune responses (Supplementary Figure 9, A & B), whereas S-B8 displayed weak to moderate autoreactivity (Supplementary Figure 9C). We further generated an S-B8 putative germline antibody, by mutating back all of the SHMs in the S-B8 heavy chain to the naïve IGHV3-66 sequence. The mutated antibody showed greater autoreactivity than S-B8 (Supplementary Figure 9D) and no S-RBD binding up to 12.5 nM (Supplementary Figure 10).

## DISCUSSION

For over a century, serology has been used to document the origin and presence of infectious agents in patients. Classically, serology depends on the actual presence of specific antibody proteins in the blood. Their target is thought to be the infectious agent, and their presence indicates a relatively recent exposure. By contrast, antibody libraries are nucleic acid based, and include genetic material from memory cells. As such, they provide a record of all of the antibodies that an individual has made, irrespective of whether they are currently being produced. This “fossil record” enabled us to discover SARS-CoV-2 neutralizing antibodies induced either by previous infection or from other immune responses. Furthermore, combinatorial antibody libraries typically yield more diverse antibodies with the desired specificity. Here, the presence of many somatic mutations in the antibodies indicates a sustained drive of the immune response to continued presence of a foreign antigen, as occurs during virus replication. Thus, the modern serology detailed here suggests that one of the individuals from whom the library was generated could have been exposed to SARS-CoV-2 or a similar virus.

Although the antibody library used here was established in 1999 before the SARS and COVID-19 pandemics (*23*), three potent neutralizing antibodies were discovered in this library. Wec et al. (*4*) identified several S-RBD-directed antibodies that potently cross-neutralize SARS-CoV (IC_50_: 0.004-0.06 μg/mL to pseudovirus) and SARS-CoV-2 (IC_50_: 0.05-1.4 μg/mL to pseudovirus) from memory B cells of a SARS donor. They also found over 80% of the low affinity SARS and SARS-2 cross-reactive antibodies reacted with one or more of the human coronavirus spike proteins, such as HCoV-NL63, HCoV-229E, HCoV-OC43, etc., indicating SARS-CoV infection may have boosted a pre-existing memory B cell response induced by circulating HCoVs (*4*). Interestingly, the three antibodies, S-B8, S-E6 and S-D4, identified in this study do not cross-react with the SARS spike protein. Moreover, autoreactivity assay in a HEp-2 cell ruled out that S-E6 and S-D4 originate from autoimmune responses, whereas S-B8 showed weak to moderate autoreactivity (Supplementary Figure 9, A – C), which was increased in the S-B8 putative germline antibody (Supplementary Figure 9D).

Our structural studies on S-E6 and S-B8 revealed several striking features of these combinatorial antibodies. The primary immune response to viral infection is followed by a secondary response that generates functionally better antibodies, where the binding energy can be refined by somatic hypermutation (*18*). The secondary immune response is for later encounter of the same antigen, and is the basis of vaccination. In cases of pandemics, such as SARS-CoV-2, avian influenza or Ebola virus, if the infection is not dealt with by the immune system in the first few days, the patient has a high probability of dying, and as a consequence, the immune system will not have enough time to refine the immune response (*35*). Consistently, neutralizing antibodies isolated from SARS-CoV-2 convalescent patients contain only a few amino-acid mutations (*3, 8, 9, 36, 37*). that may be a result of weak B cell stimulation due to rapid viral clearance. Neutralizing antibodies from convalescent patients may then possibly not be fully refined (matured) (*36*). In comparison, S-B8 and S-E6 exhibited much higher SHM, many of which are involved in specific interactions with SARS-CoV-2 RBD (S-RBD). Nine of 13 SHM residues in S-B8 and eight of 22 in S-E6 are located in the antibody-antigen interface (Supplementary Figure 6). While some of these SHM residues only use their peptide backbone, others rely on specific side chains for S-RBD binding (Figure 6, C, D, E & G). Interestingly, SHM in CDRH1 of S-E6 generates a _33_NY_34_ sequence that is similar to the 32NY33 motif in IGHV3-53/3-66 antibodies, which are the most frequent germlines used in targeting the S-RBD, indicative that the combinatorial antibody library and the maturation process can yield alternate antibody solutions (Supplementary Figure 6). However, it is unclear how these SHM residues could have been raised specifically to the SARS-CoV-2 S-RBD, since the library was generated long before the SARS-CoV-2 pandemic. Thus, these findings raise fascinating questions about the original antigen(s) that elicited S-B8 and S-E6.

The unnaturally paired antibodies in a combinatorial scFv antibody library also allow one to identify other alternative solutions with high binding affinity and efficacy (*18*). Rare germline antibodies can then potentially be enriched during the iterative affinity panning, as for example for S-E6, which is an IGHV4-31 antibody rarely seen in neutralizing antibodies from convalescent patients (Supplementary Figure 11).

A general question posed by these studies is whether therapies based on the vast number of starting antibodies in combinatorial libraries could be more powerful than the antibodies generated *in vivo* during the limited time available for affinity-based antibody evolution and selection in the setting of an acute and potentially lethal infection. This also has direct relevance in the clinical setting, where a major concern in antibody therapy is the high mutation rate of viral spike proteins, which can render prior highly specific antibodies unable to recognize or neutralize mutant viruses. Mixtures of antibodies targeting distinct epitopes can be used to overcome such immune escape (*38*). Antibodies identified from combinatorial libraries with high SHM and rare germline derivation, in combination with antibodies from convalescent patients, or with convalescent plasma, could provide yet another therapeutic option and a potential antidote to immune escape. Knowledge of the evolution of immune response in terms of the interactions of neutralizing antibodies and their binding epitopes on SARS-CoV-2 also provide a blueprint for next-generation vaccine design.

Since a complete antibody repertoire of an individual is the starting reservoir of all possibilities, one can also learn much more about the origins and evolution of an immune response against viral challenge when it is studied (*18*). The observation of highly potent neutralizing antibodies from a library of “healthy” donors before the COVID-19 pandemic could also indicate possible prior exposure of a donor(s) to a similar coronavirus 20 years ago. However, whether these antibodies are a consequence of background immunity remains to be elucidated. Notwithstanding, we are not unaware of the potential implications of such findings from libraries made decades ago concerning the origin of the virus currently circulating.

## MATERIALS AND METHODS

### Cell culture

The Vero cell line (ATCC^®^ CCL-81™) was maintained in a DMEM/F-12k media (Gibco, #C11330500CP) containing 10% (v/v) FBS (Gibco, #1600074). The FreeStyle™ 293-F (HEK 293F, ThermoFisher Scientific, #R79007) cell line was cultured in a Freestyle 293 expression media (ThermoFisher Scientific, #12338026). For establishing the HEK293T/*h*ACE2 stable cell line, HEK293T cells (ATCC^®^ ACS-4500™) were transiently transfected with *h*ACE2 fusion BFP encoding PB510 plasmid using PiggyBac Transposon System (System Biosciences, PB210PA-1), followed by addition of 2 μg/mL puromycin 6 h post-transfection. The resulting cells were kept in puromycin-containing media for an extra 2 days. Positive cells with BFP expression were sorted by a flow cytometry instrument (BD FACS Aria III). The sorted cells with overexpressed *h*ACE2 were expanded and cultured in a DMEM media (Gibco, #10566016) supplemented with 10% FBS (v/v) and 10 μg/mL puromycin.

### Expression and purification of recombinant SARS-CoV-2 spike RBD, human ACE2 and antibodies

The DNA sequences of codon-optimized SARS-CoV-2 spike Receptor Binding Domain (S-RBD) and human ACE2 extracellular domain (*h*ACE2-ECD) were cloned into a *p*Fuse-Fc expression vector (Invivogen). A thrombin cleavage sequence was inserted between RBD and Fc to generate a cleavable human Fc tag for future studies. The SARS-CoV-2 S-RBD-*h*Fc and *h*ACE2-ECD-*m*Fc proteins were heterologously expressed in HEK293F cells by transient transfection and cultured for 4 days, then purified by Mabselect columns (Cytiva, #17-5199-01). Briefly, cell media with secreted Fc tagged recombinant proteins, S-RBD-*h*Fc and *h*ACE2-ECD-*m*Fc, were loaded onto a Mabselect column that was pre-washed and equilibrated with a PBS buffer (150 mM NaCl, 20 mM sodium phosphate, pH 7.2), and eluted using a pH 3.4 citrate acid buffer.

DNA sequences for the variable regions of the combinatorial antibodies were cloned into a full-length human IgG4 mutant construct (S228P) and expressed in HEK293F cells for 4 days and further purified by Mabselect chromatography. Purified recombinant proteins and antibodies were buffer-exchanged into a PBS buffer using centrifugal concentrators.

### Function-guided phage panning

SARS-CoV-2 S-RBD specific scFv antibodies were selected from a combinatorial human monoclonal scFv antibody phage library (10^11^ members) after two rounds of affinity enrichment against the biotinylated S-RBD protein immobilized on the streptavidin-coated magnetic beads (Pierce, #21925), followed by a third round of competitive panning *vs*. *h*ACE2-ECD protein. Briefly, phagemid (displaying the antibody library) binding to the antigen (S-RBD) was enriched at each cycle and eluted with Glycine-HCl (pH 2.2) in the first two rounds of screening. XL1-Blue cells were used to express and amplify the output phagemids for the next round of panning. To determine *h*ACE2 competitive antibodies, a kinetic competitive panning method was adopted in the third round panning. Instead of the conventional pH 2.2 buffer, an elution buffer containing a saturated concentration of *h*ACE2-ECD protein (200 nM; for S-RBD and *h*ACE2-ECD binding, EC_80_ = 80 nM) was used to elute the phagemids twice. After three iterations, 96 positive colonies were selected and analyzed by phage ELISA as described (*39*). All of the positive clones were sequenced using Sanger sequencing. Both the DNA and protein sequences of CDR3 domains were analyzed using the international ImMunoGeneTics (IMGT) information platform (http://www.imgt.org/).

### Phage ELISA

Avidin (Pierce, #21121) was diluted to a final concentration of 2 ng/μL in a PBS buffer (Sigma, #C3041). The resulting avidin solution was used to coat the 96-half well plates (25 μL/well) at 4 °C overnight. The coated plates were washed once with the PBS buffer (150 μL/well) followed by the addition and incubation of 25 μL biotinylated SARS-CoV-2 S-RBD-*h*Fc solution (2 ng/μL) in each well at room temperature for 1 h. The PBST (PBS containing 0.05% Tween-20) buffer alone and the *h*Fc solution (2 ng/μL) were used as the background and negative controls, respectively. After removal of the incubation solution, the resulting plates were rinsed once using the PBST buffer and incubated with a blocking solution containing 5% milk (v/v) in PBST (150 μL/well) at 37 °C for 1 h. After blocking and PBST washing (once), 50 μL of phagemid-containing XL1-Blue culture medium supernatants (by centrifuging the third round panning output XL1-Blue cells at 3000 g, 15 min) mixed with 10 μL 5% milk (v/v) in PBST was added to each well and incubated at 37 °C for 1 h. The resulting plates were rinsed eight times using PBST before subjecting to horseradish peroxidase (HRP) detection. A solution containing the secondary antibody, anti-M13 bacteriophage antibody conjugated with HRP (dilution factor 1 : 5000; Sino Biological, #11973-MM05T-H), was added into the above plates (150 μL/well) and incubated at 37 °C for 1 h. Plates were then washed eight times with PBST followed by the addition of 50 μL ABTS solution (Roche, #11684302001) into each well. After ~10 min incubation at room temperature, the absorbance change at 405 nm in each well was measured on a microplate reader (Enspire, PerkinElmer).

### Competitive ELISA

Competition between the selected antibodies and *h*ACE2 for binding to the SARS-CoV-2 spike protein RBD was measured. The recombinant *h*ACE2-ECD was coated in PBS buffer at 2 ng/μL, 100 μL per well at 4 °C overnight, washed with PBS once, then blocked with 3% BSA in PBS. Biotinylated S-RBD (*h*Fc tag removed by thrombin digestion) at a final concentration of 50 nM was incubated with 2-fold serial diluted S-B8, S-D4, and S-E6 antibodies (from 1 - 133 nM) at 4 °C for 30 min, in which an IgG4e1 isotype antibody was used as the negative control. The S-RBD and antibody mixture was then added to the *h*ACE2-ECD coated plates and incubated at room temperature for 1 h, followed by 4 washes with PBST. The *h*ACE2-ECD bound S-RBD in the plate was detected using a Streptavidin-HRP conjugated protein.

### Affinity determination by Biolayer Interferometry (BLI)

Binding affinities of S-D4 with SARS-CoV-2 S-RBD were performed by BLI on an Octet RED96 (Molecular Devices LLC, San Jose, CA, USA) using AR2G biosensors. The SARS-CoV-2 S-RBD fused *h*Fc was first digested by thrombin to remove the Fc tag. The resulting S-RBD diluted in a PBS solution containing 0.02% Tween-20 and 0.05% BSA (PBST-B) (10 μg/mL) was loaded to the AR2G biosensor by amine coupling. The AR2G-S-RBD sensors were dipped into a PBST-B for 60 sec to establish a baseline, and then incubated with 2-fold serial diluted antibody solutions to record the progressive curves of association. Finally, sensors were incubated in a PBST-B buffer to record the progressive curves of dissociation. For S-B8 and S-E6 detections, S-RBD was first biotinylated before loading to a streptavidin (SA) sensor, the remaining procedure was same to that of S-D4. Sensor regeneration was performed by dipping the used sensors into a pH 3.4 citrate acid buffer, and equilibrated in a PBST-B buffer. Results were analyzed by ForteBio Data Analysis software.

### Interaction of antibodies with cell surface expressed spike by FACS

In a flow-cytometry binding experiment, the spike protein of either full-length SARS-CoV-2 or SARS, which was conjugated with P2A-EGFP, was transiently transfected into a HEK293T cell. After 24 h cultivation, cells were collected and re-suspended in an ice-cold FACS buffer (PBS, 0.05 % BSA and 2 mM EDTA). The spike protein expressing cells (50000 cells per tube) were then incubated with different anti-S-RBD antibodies for 20 min at 4 °C, and washed with 1 mL ice-cold FACS buffer, spun, and re-suspended in a 100 μL ice-cold FACS buffer containing the Alexa555 conjugated secondary antibody that recognizes human Fc (1 : 800 v/v dilution, Life technology, # A21433). After incubating at 4 °C for 15 min, the cells were washed twice and re-suspended in a FACS buffer, and then sorted and analyzed on a flow cytometer (CytoFLEX S, Beckman Culter) to determine relative binding level by the antibodies to the cell overexpressing wild-type spikes. Mean fluorescence intensities of Alexa555 in eGFP-positive cells were recorded and analyzed to evaluate antibody binding.

### Size-exclusion-high-performance liquid chromatography (SEC-HPLC)

Twenty μL of 0.5 μg/μL purified S-RBD antibodies were applied to an Agilent Bio SEC-5, 500A HPLC system. The mobile phase used PBS buffer (pH 7.2) running at a flow rate of 0.35 mL/min. Absorbance was analyzed and integrated by retention time and area under the curve (AUC) to determine the percentage of aggregation, monomer and degradants compositions.

### Cell-cell fusion assay

The cell-cell fusion assay was established according to a previous report (*40*) with minor modifications. Briefly, *h*ACE2 positive Vero cells (cells with endogenous *h*ACE2 were sorted by FACS) were used as target cells. HEK293F cells that are transiently transfected with either SARS-CoV-2 spike-P2A-EGFP or SARS spike-P2A-EGFP were set as effector cells. The target Vero cells were first seeded into 24-well plates at a density of 1×10^5^/well and cultivated at 37 °C for 4 h, followed by addition of effector cells, HEK293F/SARS spike-EGFP or HEK293F/SARS-CoV-2 spike-EGFP, at a ratio of 2:1, respectively. The co-cultures of cells were cultivated in a DMEM medium with 10% FBS, and treated with or without anti-SARS-CoV-2 spike antibodies at indicated concentrations. The recombinant SARS-CoV-2 S-RBD was used as a positive control. After cultivating at 37 °C for 6 h, the rates of cell-cell fusion were evaluated using a fluorescence microscope (EVOS M5000, Life Technologies). Five fields for microscopic analysis were randomly selected in each treated group, the numbers of fused and unfused EGFP positive cells were counted.

### Preparation of pseudovirus

HEK293T cells were co-transfected with both NL4-3 mCherry Luciferase plasmid (addgene #44965) and pcDNA3.1 SARS-CoV-2 spikeΔ19 plasmid (encoding SARS-CoV-2 spike protein, with 19 AA truncated in C terminal) using Lipofectamine 3000 (Invitrogen, L3000-015) following the manufacturer’s instruction. Pseudotyped particles were readily released into the supernatant. The supernatants containing SARS-CoV-2 pseudovirus were harvested at 48 h post-transfection, filtered (0.45 μm pore size, Sartorius, 16533-K), and mixed with the Lenti-X Concentrator (Takara, 631231) overnight at 4 °C. The mixture was then centrifuged at 1500 g for 45 min at 4 °C. The cell pellets were collected and re-suspended in a DMEM medium and stored at −80 °C until use.

### Pseudovirus-based neutralization assay

To detect the neutralization ability of selected antibodies against infection of coronavirus pseudovirus (PSV), HEK293T/*h*ACE2 cells were first seeded into 96-well white bottom plates at a density of 1×10^4^/well, and cultivated overnight. The PSV was pre-incubated with an equal volume of different concentrations of selected antibodies (dilution factor: 3.16, from 200 nM to 200 fM for S-B8 and S-D4, 200 nM to 6.3 fM for S-E6) in DMEM at 37 °C for 30 min. DMEM with or without PSV in the absence of antibodies were set as controls. After incubation, the PSV mixture was transferred to the culture plates containing HEK293T/*h*ACE2 cells. The DMEM media containing PSV and antibodies were replaced with fresh media after 16 h treatment, cells were incubated for an additional 48 h. PSV infection efficacy was evaluated by luciferase activity using Bright-Lumi™ Firefly Luciferase Reporter Gene Assay Kit (Beyotime, RG015M). Fifty microliter of luciferase substrate was added to each well, and the relative luminescence unit (RLU) values were measured on an Envision plate reader (PerkinElmer, Ensight).

### Authentic SARS-CoV-2 virus neutralization assay

The study was performed in the CL3 Facility of the University of Oxford operating under license from the HSE, on the basis of an agreed Code of Practice, Risk Assessments (under ACDP) and Standard Operating Procedures. A detailed protocol with supporting data will be prepared shortly (Harding, A, Gilbert-Jaramillo, J, et al., 2020). In brief, this rapid, high-throughput assay determines the concentration of antibody that produces a 50% reduction in infectious focus-forming units of authentic SARS-CoV-2 in Vero cells, as follows. Quadruplicate, 0.5log10 serial dilutions of antibody (11 steps from 100 nM to 1 pM) were pre-incubated with a fixed dose of SARS-CoV-2 (Victoria 01/2020 isolate) before incubation with Vero cells. A 1.5% carboxymethyl cellulose-containing overlay was used to prevent satellite focus formation. Twenty hours post-infection, the monolayers were fixed with 4% paraformaldehyde, permeabilized with 2% Triton X-100 and stained for N antigen using mAb EY 2A (*41*). After development with a peroxidase-conjugated antibody and True Blue peroxidase substrate, infectious foci were enumerated by ELISPOT reader. Data were analyzed using four-parameter logistic regression (Hill equation) in GraphPad Prism 8.3.

### Autoreactivity assay

The autoreactivity assay was performed using a HEp-2 anti-nuclear antibodies (ANA) kit (Medical & Biological Laboratories Co., Ltd, #4220-12CN) according to the manufacturer’s instructions. Briefly, 35 μL of 0.1 mg/mL antibodies were loaded to the wells in a slide pre-seeded with fixed and permeabilized HEp-2 cells and incubated for 20 min at room temperature. Positive serum from autoimmune patients and negative serum from healthy donors provided by the kit were used as controls. After washing twice (5 min each), the FITC-conjugated secondary anti-human antibody was incubated with the cells for 20 min at room temperature. The slide was then washed and mounted with a coverslip before observation on a fluorescent microscope (ZEISS, Axio Observer A1) with a 20× objective.

### Protein production and structure determination

The coding sequence for receptor binding domain (RBD; residues 319-541) of the SARS-CoV-2 spike (S) protein was synthesized and cloned into a customized pFastBac vector (*42*), which was designed to fuse an N-terminal gp67 signal peptide and C-terminal His6-tag to the target protein. To express the RBD protein, a recombinant bacmid DNA was generated from the sequencing-confirmed pFastBac construct using the Bac-to-Bac system (Life Technologies). Baculovirus was generated by transfecting purified bacmid DNA into Sf9 cells using FuGENE HD (Promega), and subsequently used to infect suspension cultures of High Five cells (Life Technologies) at a multiplicity of infection (MOI) of 5 to 10. Infected High Five cells were incubated at 28 °C with shaking at 110 rpm for 72 h for protein expression. RBD protein that was secreted into the supernatant, harvested, and then concentrated with a 10 kDa MW cutoff Centramate cassette (Pall Corporation). The RBD protein was purified by affinity chromatography using Ni-NTA resin (QIAGEN), followed by size exclusion chromatography on a HiLoad Superdex 200 pg column (GE Healthcare), and buffer exchanged into 20 mM Tris-HCl pH 7.4 and 150 mM NaCl using the same protocol as previously described (*43*). Fabs were expressed in ExpiCHO cells and purified using CaptureSelect CH1-XL resin (ThermoFisher) and followed by size exclusion chromatography. The Fab/RBD complexes were formed by mixing the two components in an equimolar ratio and incubating overnight at 4 °C before setting-up crystal trials. The Fab/RBD complexes were screened for crystallization using 384 conditions of the JCSG Core Suite (QIAGEN) on our robotic CrystalMation system (Rigaku) at The Scripps Research Institute. Crystals appeared in the first week, were harvested during the second week, and then flash-cooled in liquid nitrogen for X-ray diffraction experiments. Diffraction data were collected at cryogenic temperature (100 K) at beamline 23-ID-B of the Advanced Photon Source (APS) at Argonne National Laboratory with a beam wavelength of 1.033 Å and processed with HKL2000 (*44*). Diffraction data were collected from crystals grown in conditions: 20% PEG 3350, 0.2 M sodium sulfate, pH 6.6 for S-B8/RBD complex; 20% isopropanol, 20% PEG 4000, 0.1 M citrate pH 5.6 for S-E6/RBD complex. The X-ray structures were solved by molecular replacement (MR) using PHASER (*45*) with MR models for the RBD and Fab from PDB 7JMW (*46*). Iterative model building and refinement were carried out in COOT (*47*) and PHENIX (*48*), respectively. Epitope and paratope residues, as well as their interactions, were identified by using PISA program (*49*) with buried surface area (BSA >0 Å^2^) as the criterion.

### Data analysis and statistics

The results were expressed as means ± standard deviation (SD) unless otherwise indicated. Data analysis was performed by one-way analysis of variance (ANOVA) using Origin Pro 2019 statistical software or GraphPad Prism software. Significance was assumed at a *P* value < 0.05.

## ACKNOWLEDGEMENTS

We thank Dr. Lichun Jiang, Dr. Wei Wang and Zhangyue Song from the Biomedical Big Data platform of the Shanghai Institute for Advanced Immunochemical Studies (SIAIS) of ShanghaiTech University for sequencing and data analysis, Pengwei Zhang, Dr. Lishuang Zhang and Juan Kong from the high-throughput screening platform of SIAIS for technical support in cell sorting and phage panning, Jiakang Chen from the Analytical Chemistry Platform of SIAIS for technical support in SEC-HPLC.

## AUTHOR CONTRIBUTIONS

R.A.L., G.Y., H.W., W.J., R.A.D, and I.A.W. conceived the project. M.Q., P.X.M., P.D.T., and X.J.S. contributed to project design and extensive discussions. M.Q., P.X.M., Y.L., L.L.L., C.Y.M., Q.J., P.D.T., F.L.W, Z.A.L., A.S. and Y.Z. performed antibody selection, identification, binding, cell-cell fusion and pseudovirus neutralization work. H.J.L., M.Y., N.C.W., C-C.D.L., X.Y.Z. performed structural work involving protein production, crystallization, structure determination and analysis. W.J., A.H. and J.G-J. performed the authentic virus neutralization experiments. G.Y., M.Q., P.X.M., I.A.W. and R.A.L. analyzed data and wrote the manuscript. X.X.H., W.J. and R.A.D. provided manuscript edits and suggestions.

## DATA AND MATERIALS AVAILABILITY

Correspondence and requests for materials should be addressed to Richard A. Lerner. X-ray coordinates and structure factors for S-B8 and S-E6 in complex with the SARS-CoV-2 RBD have been deposited in the RCSB Protein Data Bank under accession codes: 7KN3, 7KN4.

---

**Supplementary Figure 1.**
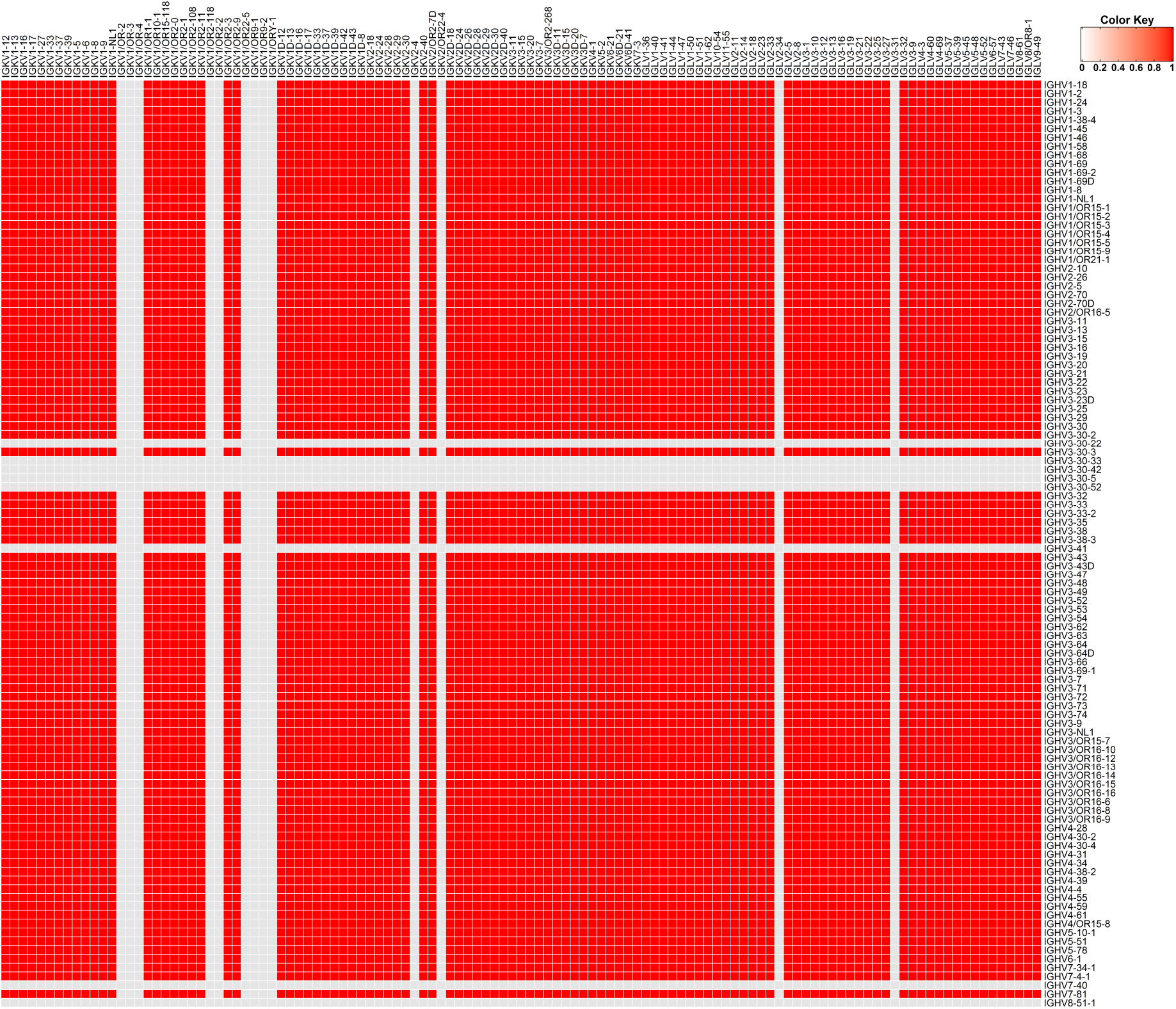
Germline matrix coverage of the naïve antibody library. The heavy and light chain genes of the naïve antibody library were separately sequenced by next generation sequencing, and the IGHV *vs*. IGLV and IGKV germline matrix coverage hotmaped (red color) are shown aligned to the full germline distribution derived from IMGT (international ImMunoGeneTics) database (http://www.imgt.org/).

**Supplementary Figure 2.**
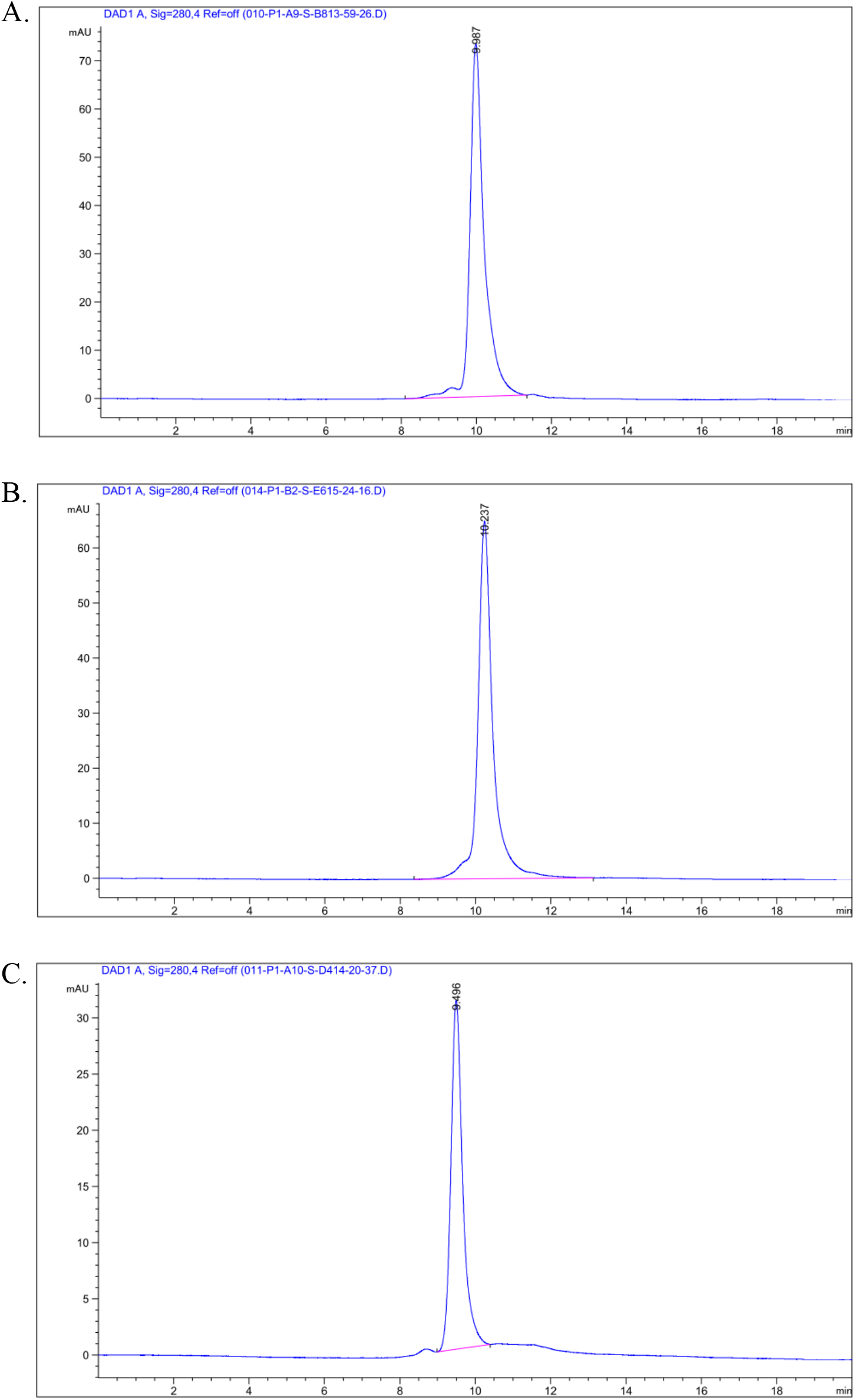
SEC-HPLC of selected antibodies. SEC-HPLC analysis of S-B8 (**A**), S-D4 (**B**) and S-E6 (**C**) in full length IgG4e1 formats are shown. The antibody used for analysis was at 0.5 mg/mL.

**Supplementary Figure 3.**
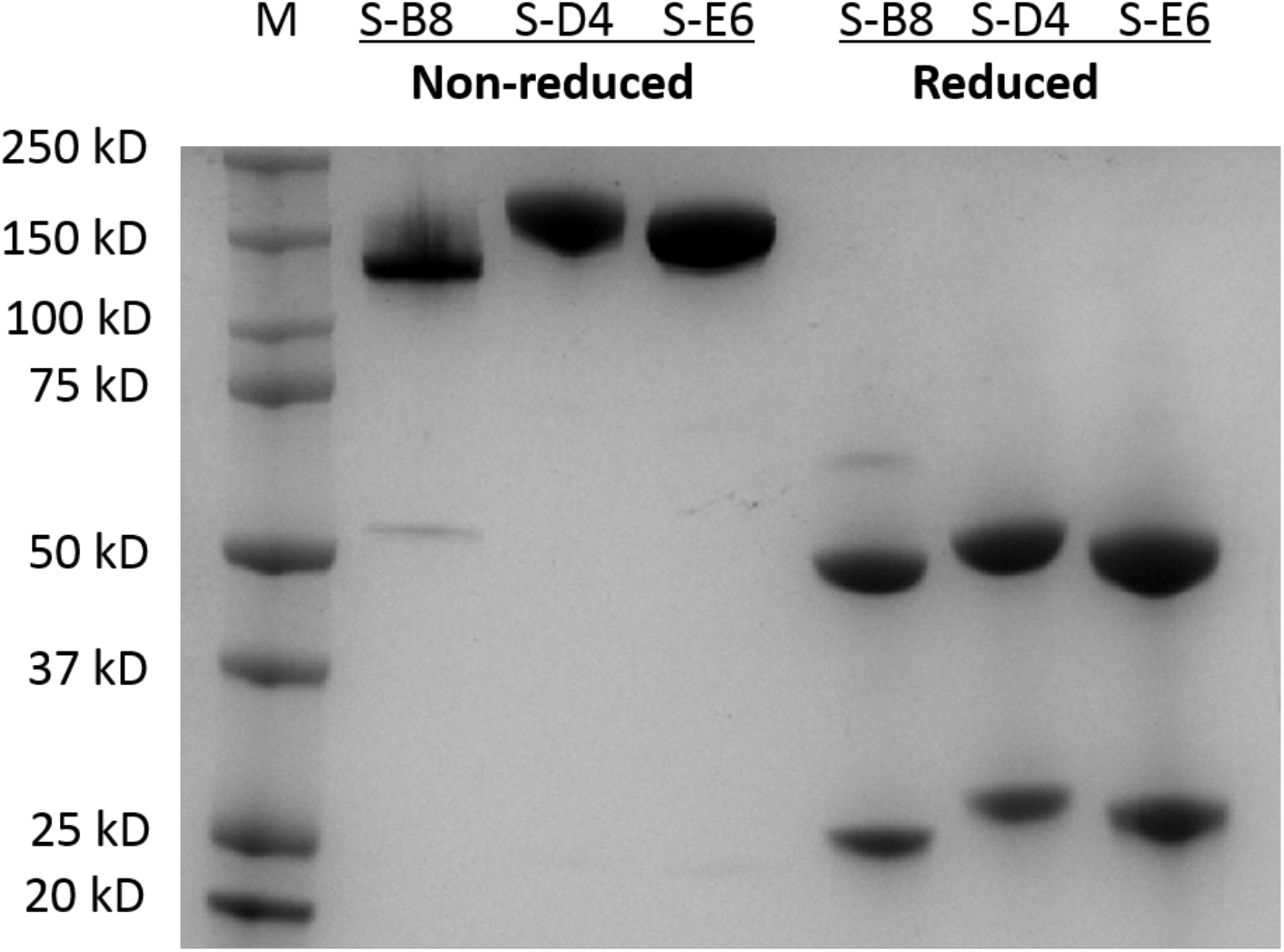
SDS-PAGE analysis of *h*ACE2 competitive antibodies. Three *h*ACE2 competitive antibodies in IgG4e1 format are analyzed by SDS-PAGE, in both non-reduced and reduced forms.

**Supplementary Figure 4.**
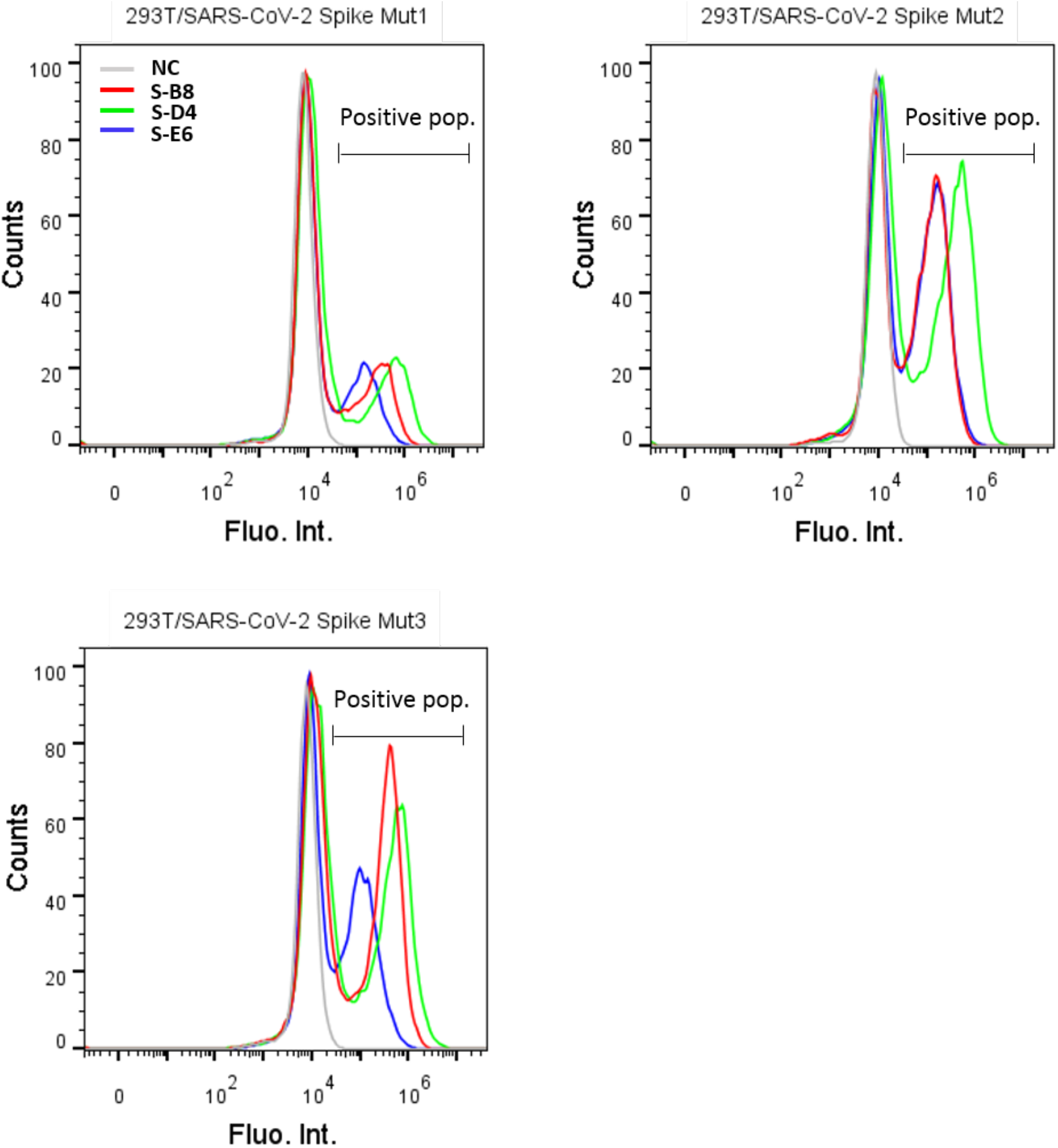
FACS analysis of antibody binding to cell surface-expressed mutated spike protein. HEK293T cells transfected with expression plasmid encoding the mutated full-length spike of SARS-CoV-2 were incubated with the three *h*ACE2 competitive IgG4 antibodies. The cells were then stained with FITC labeled anti-human IgG Fc secondary antibody and analyzed by FACS. Cells stained with only secondary antibody were set as negative control (NC). Positive binding cells populations were labeled as positive pop.

**Supplementary Figure 5.**
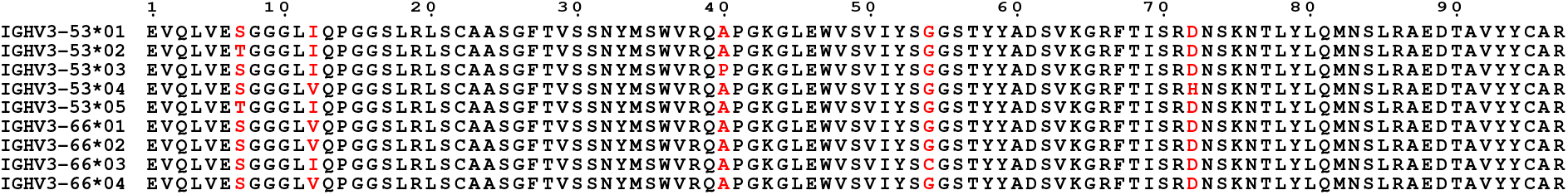
Sequence alignment of IGHV3-53 and IGHV3-66. Germline sequences from all known alleles are analyzed. Positions with any amino-acid variation are shown in red.

**Supplementary Figure 6.**
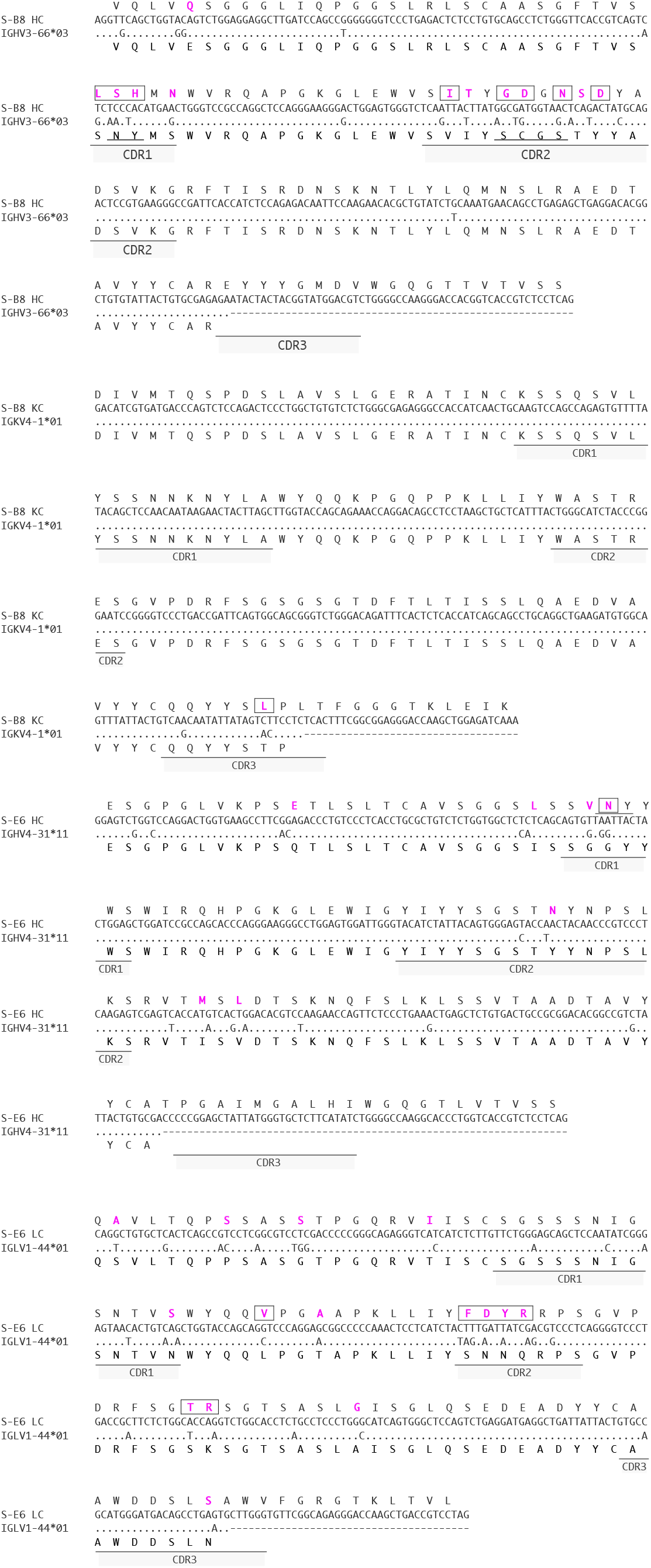
Germline sequences and somatic hypermutation (SHM) of S-B8 and S-E6. Germline sequences are aligned to heavy and light chain sequences for each antibody. Residues from SHM are colored in purple and residues interacting with SARS-CoV-2 S-RBD are boxed.

**Supplementary Figure 7.**
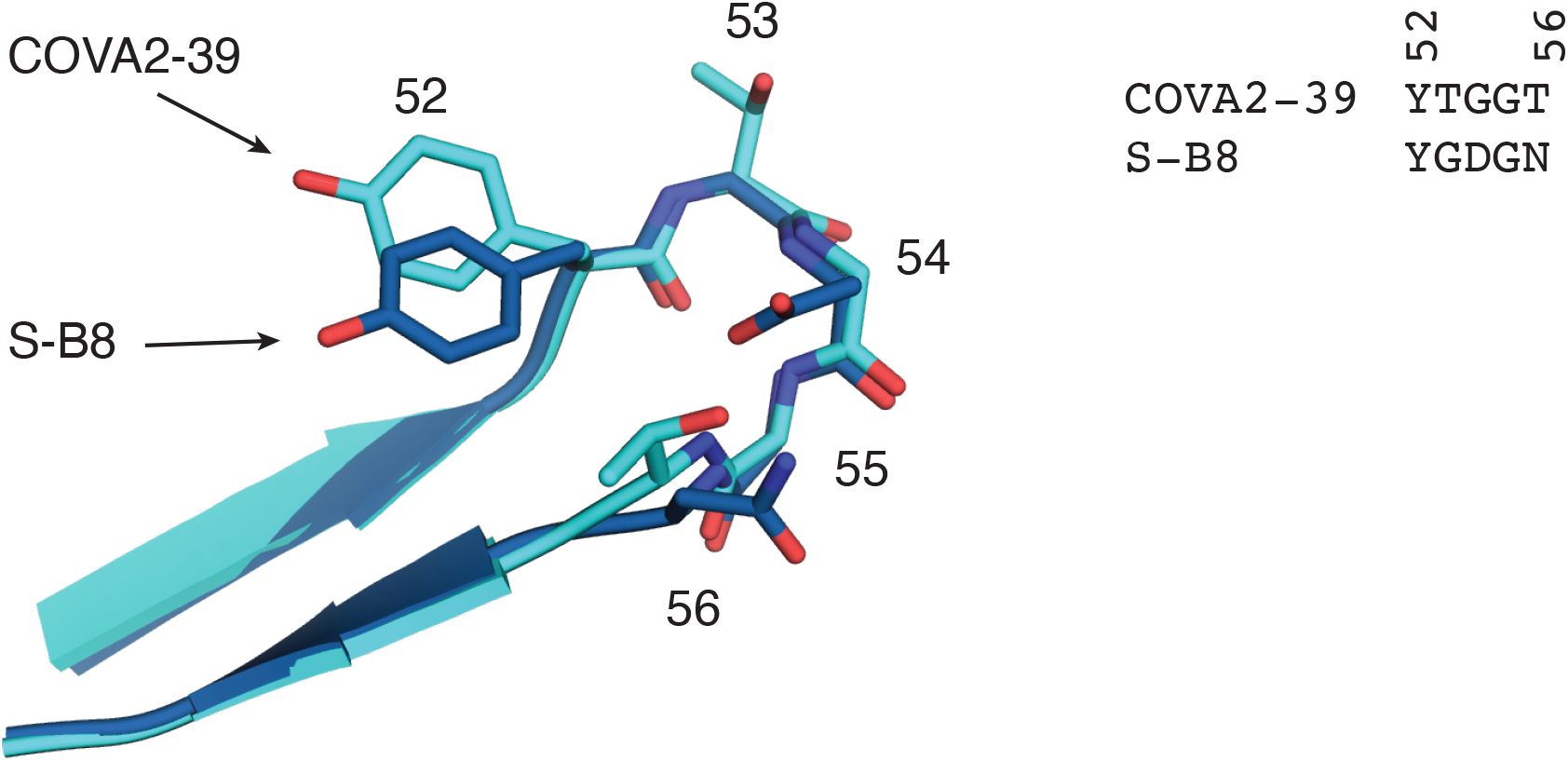
Structural comparison of CDRH2 between S-B8 and COVA2-39 (PDB 7JMP). Residues in the β-turn region are shown as sticks with corresponding sequences shown on the right and sequence numbers labeled.

**Supplementary Figure 8.**
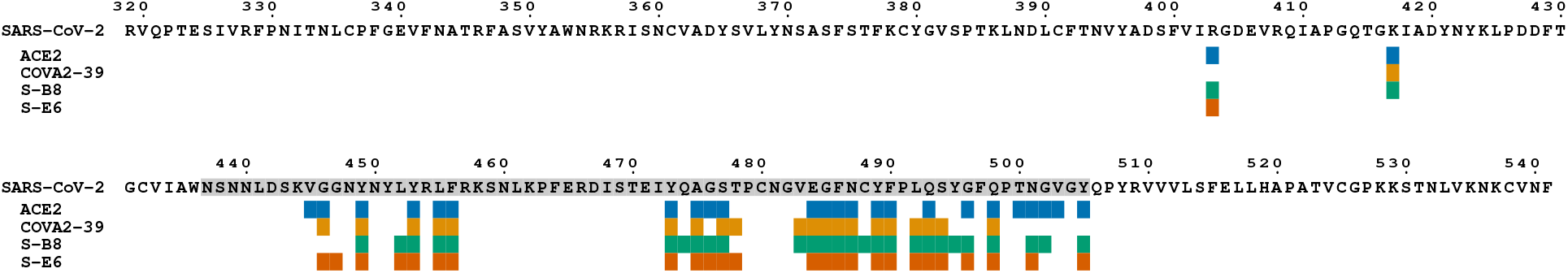
Fab epitopes and *h*ACE2 binding site. SARS-CoV-2 S-RBD epitopes of S-B8, S-E6 and COVA2-39 (PDB 7JMP) as well as *h*ACE2 binding site on the S-RBD were analyzed with PISA program using buried surface area (BSA)>0 Å^2^ as the criterion. Each antibody epitope or *h*ACE2 binding site is shown as blocks below the S-RBD sequence under the corresponding residue position. The sequence of the receptor binding motif (RBM) is shaded in grey.

**Supplementary Figure 9.**
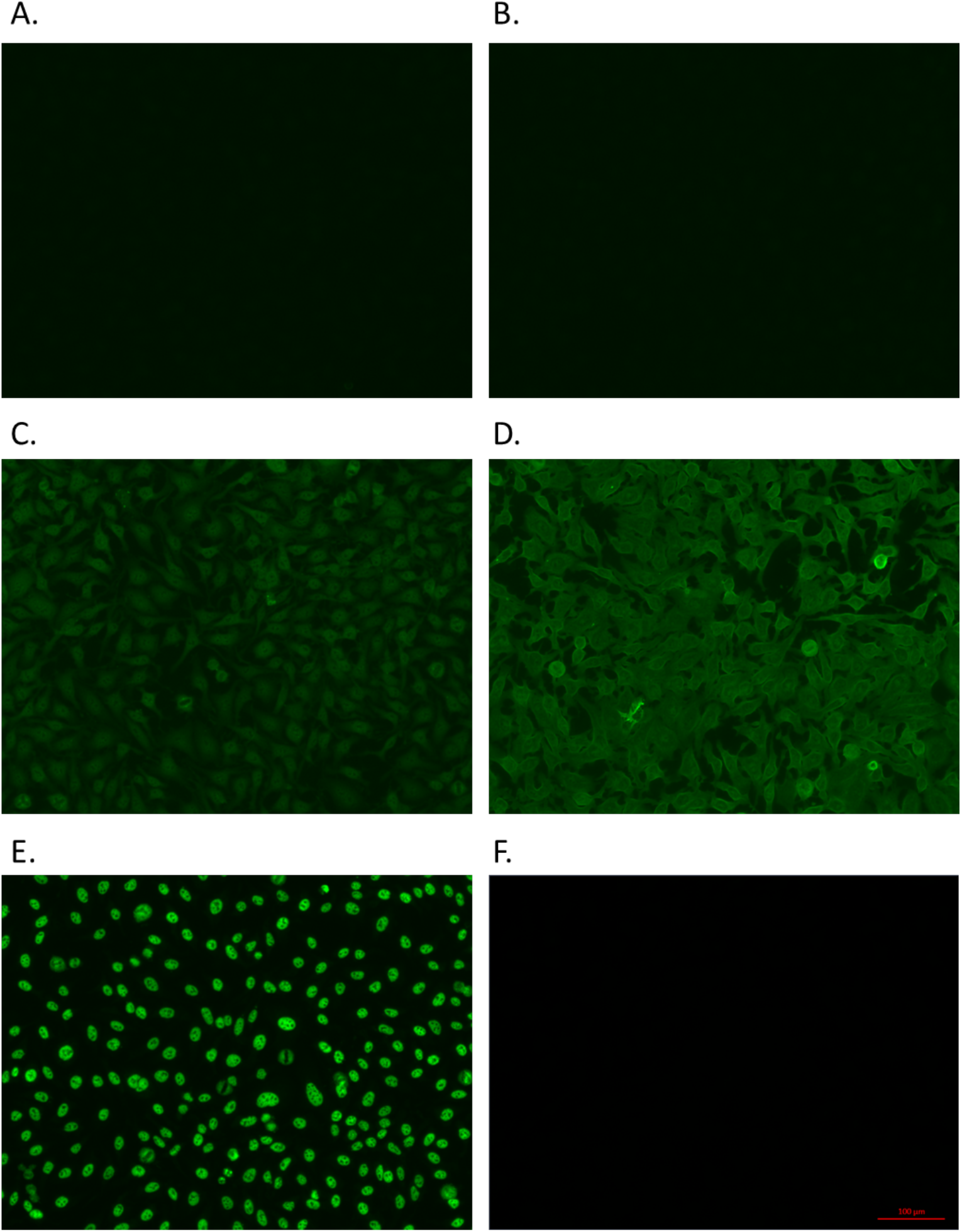
Autoreactivity detection of selected antibodies. Autoreactivity of S-D4 (**A**), S-E6 (**B**), S-B8 (**C**) and the S-B8 putative germline antibody (**D**) were tested by using a HEp-2 cell based antinuclear-antibody kit. The S-B8 putative germline antibody was generated by mutating the SHMs of S-B8 back to the germline sequence of IGHV3-66 as shown in Supplementary Figure 6. Positive control (PC) (**E**) and negative control (NC) (**F**) are from serum of patients with or without autoimmune disease, which is included in the kit. The green color indicates positive binding of the tested antibodies to HEp-2 cells. Bar=100 μm.

**Supplementary Figure 10.**
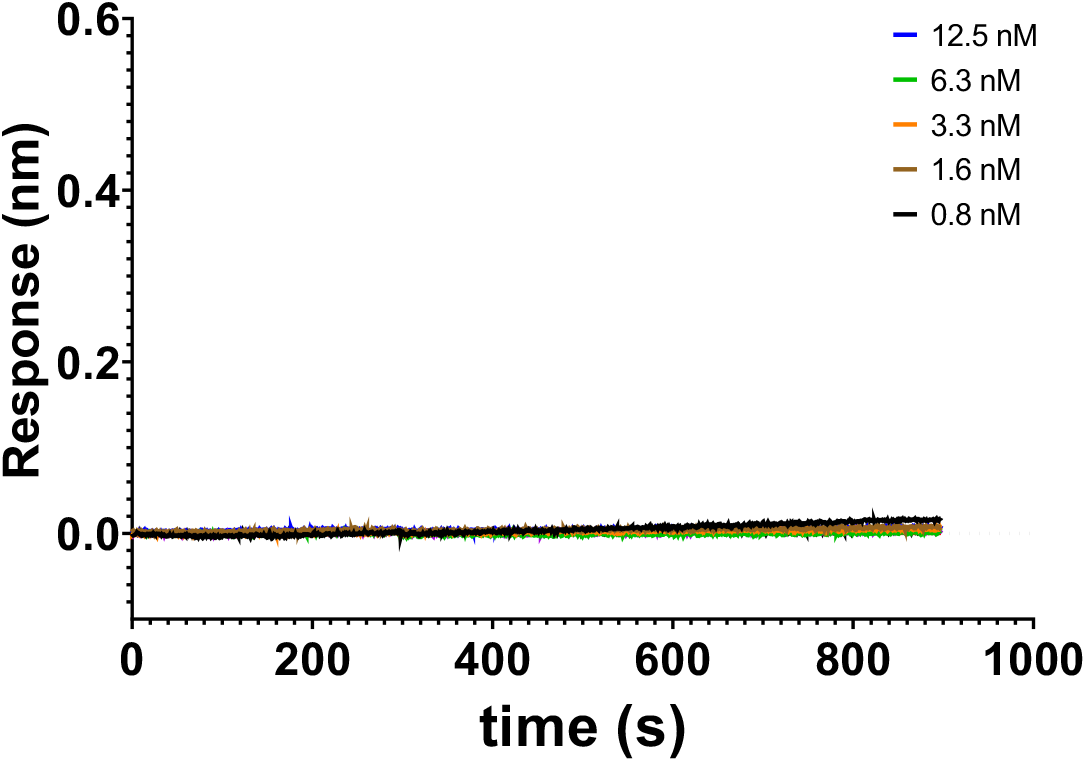
Binding kinetics of S-B8 putative germline antibody to the spike protein. Binding kinetics were measured by biolayer interferometry (BLI). Biotinylated S-RBD was loaded to the SA biosensor for detection of binding kinetics with the S-B8 putative germline antibody.

**Supplementary Figure 11.**
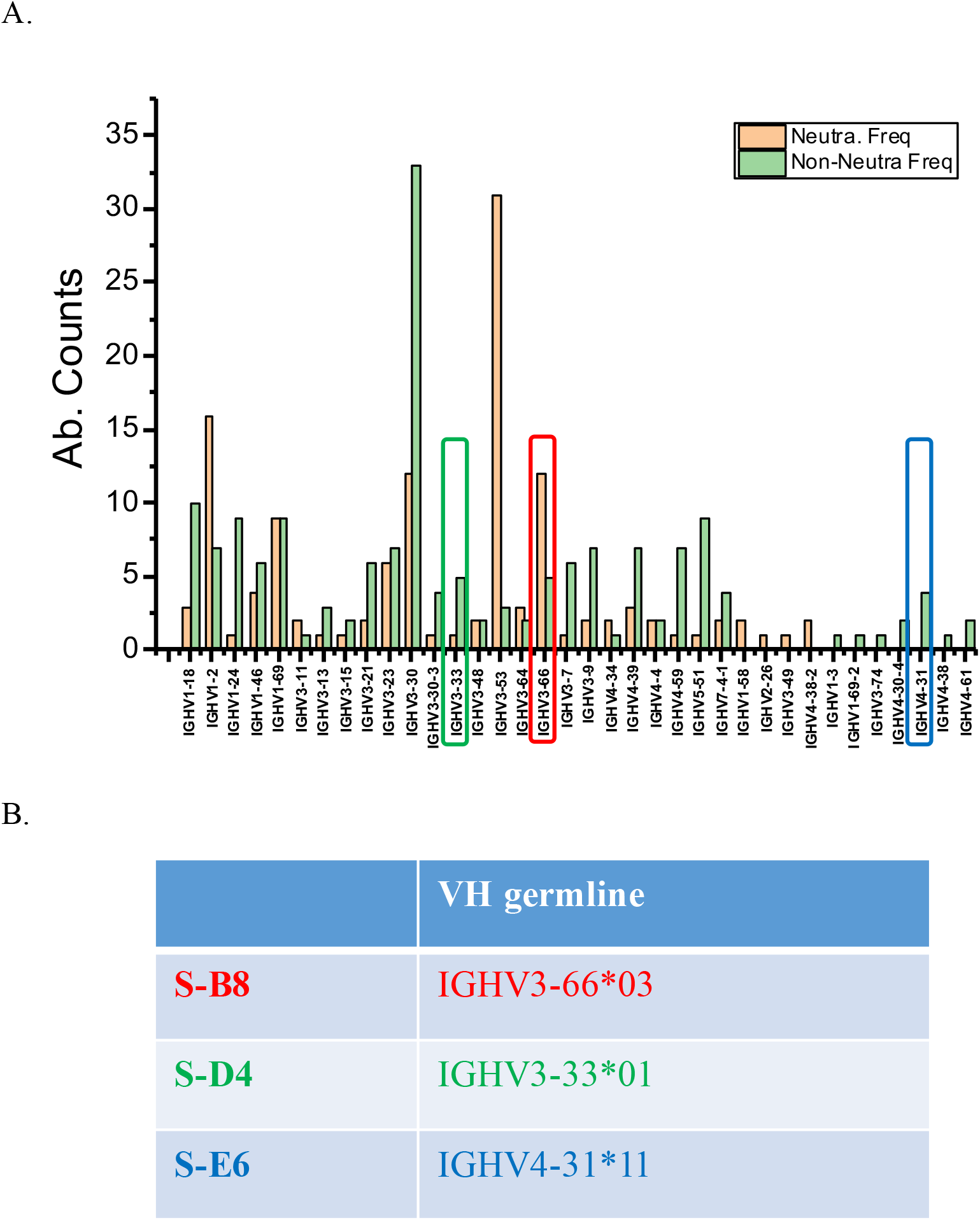
IGHV germline distribution of SARS-CoV-2 spike-targeting antibodies. (**A**) IGHV distribution of about 300 SARS-CoV-2 binding antibodies reported in the public database (http://opig.stats.ox.ac.uk/webapps/covabdab/#collapseOne) were analyzed as of June 20^th^, 2020. The numbers of neutralizing and non-neutralizing antibodies of certain germlines were also calculated and shown. (**B**) The IGHV of three competitive antibodies are indicated and shown in different colors in the Table and in **A**.

**Supplementary Table 1.** Crystallographic data collection and refinement statistics.

|  | S-B8 + RBD | S-E6 + RBD |
| --- | --- | --- |
| **Data collection** |  |  |
| Beamline | APS 23ID-B | APS 23ID-B |
| Wavelength (Å) | 1.03317 | 1.03373 |
| Space group | *R* 1 3 1 | *C* 1 2 1 |
| Unit cell parameters |  |  |
| a, b, c (Å) | 191.6, 191.6, 117.4 | 242.1, 70.2, 91.9 |
| α, β, γ (°) | 90, 90, 120 | 90, 108.5, 90 |
| Resolution (Å)<sup>a</sup> | 50.0-2.25 (2.30- | 50.0-2.70 (2.75- |
| Unique reflections<sup>a</sup> | 74,079 (3,622) | 37,826 (1,647) |
| Redundancy<sup>a</sup> | 6.4 (2.6) | 3.5 (2.6) |
| Completeness (%)<sup>a</sup> | 97.4 (71.4) | 93.3 (82.8) |
| $\langle I/\sigma \rangle$<sup>a</sup> | 17.3 (1.4) | 12.5 (1.7) |
| $R_{\text{sym}}^b$ (%)<sup>a</sup> | 9.8 (44.6) | 9.2 (56.0) |
| $R_{\text{pim}}^b$ (%)<sup>a</sup> | 4.0 (28.5) | 5.5 (38.0) |
| $CC_{1/2}^c$ (%)<sup>a</sup> | 99.4 (77.5) | 98.4 (72.0) |
| **Refinement statistics** |  |  |
| Resolution (Å) | 47.9-2.25 | 49.3-2.70 |
| Reflections (work) | 70,217 | 33,627 |
| Reflections (test) | 3,821 | 1,791 |
| $R_{\text{cryst}}^d / R_{\text{free}}^e$ (%) | 17.9/22.1 | 23.6/28.3 |
| No. of atoms | 10,193 | 8,955 |
| Macromolecules | 9,665 | 8,927 |
| Glycans | 28 | 28 |
| Solvent | 500 | - |
| Average *B*-value | 42 | 47 |
| Macromolecules | 42 | 47 |
| Fab | 40 | 43 |
| RBD | 46 | 57 |
| Glycans | 65 | 113 |
| Solvent | 45 | - |
| Wilson *B*-value (Å<sup>2</sup>) | 37 | 44 |
| **RMSD from ideal geometry** |  |  |
| Bond length (Å) | 0.005 | 0.004 |
| Bond angle (°) | 0.78 | 0.67 |
| **Ramachandran statistics (%)** |  |  |
| Favored | 97.8 | 95.5 |
| Outliers | 0.0 | 0.2 |
| **PDB code** | 7KN3 | 7KN4 |

<sup>a</sup> Numbers in parentheses refer to the highest resolution shell.
<sup>b</sup> $R_{\text{sym}} = \sum_{hkl} \sum_i |I_{hkl,i} - \langle I_{hkl} \rangle| / \sum_{hkl} \sum_i I_{hkl,i}$ and $R_{\text{pim}} = \sum_{hkl} (1/(n-1))^{1/2} \sum_i |I_{hkl,i} - \langle I_{hkl} \rangle| / \sum_{hkl} \sum_i I_{hkl,i}$ , where $I_{hkl,i}$ is the scaled intensity of the *i*th measurement of reflection *h*, *k*, *l*, $\langle I_{hkl} \rangle$ is the average intensity for that reflection, and *n* is the redundancy.
<sup>c</sup> $CC_{1/2}$ = Pearson correlation coefficient between two random half datasets.
<sup>d</sup> $R_{\text{cryst}} = \sum_{hkl} |F_o - F_c| / \sum_{hkl} |F_o| \times 100$ , where $F_o$ and $F_c$ are the observed and calculated structure factors, respectively.
<sup>e</sup> $R_{\text{free}}$ was calculated as for $R_{\text{cryst}}$ , but on a test set comprising 5% of the data excluded from refinement.

**Supplementary Table 2.**
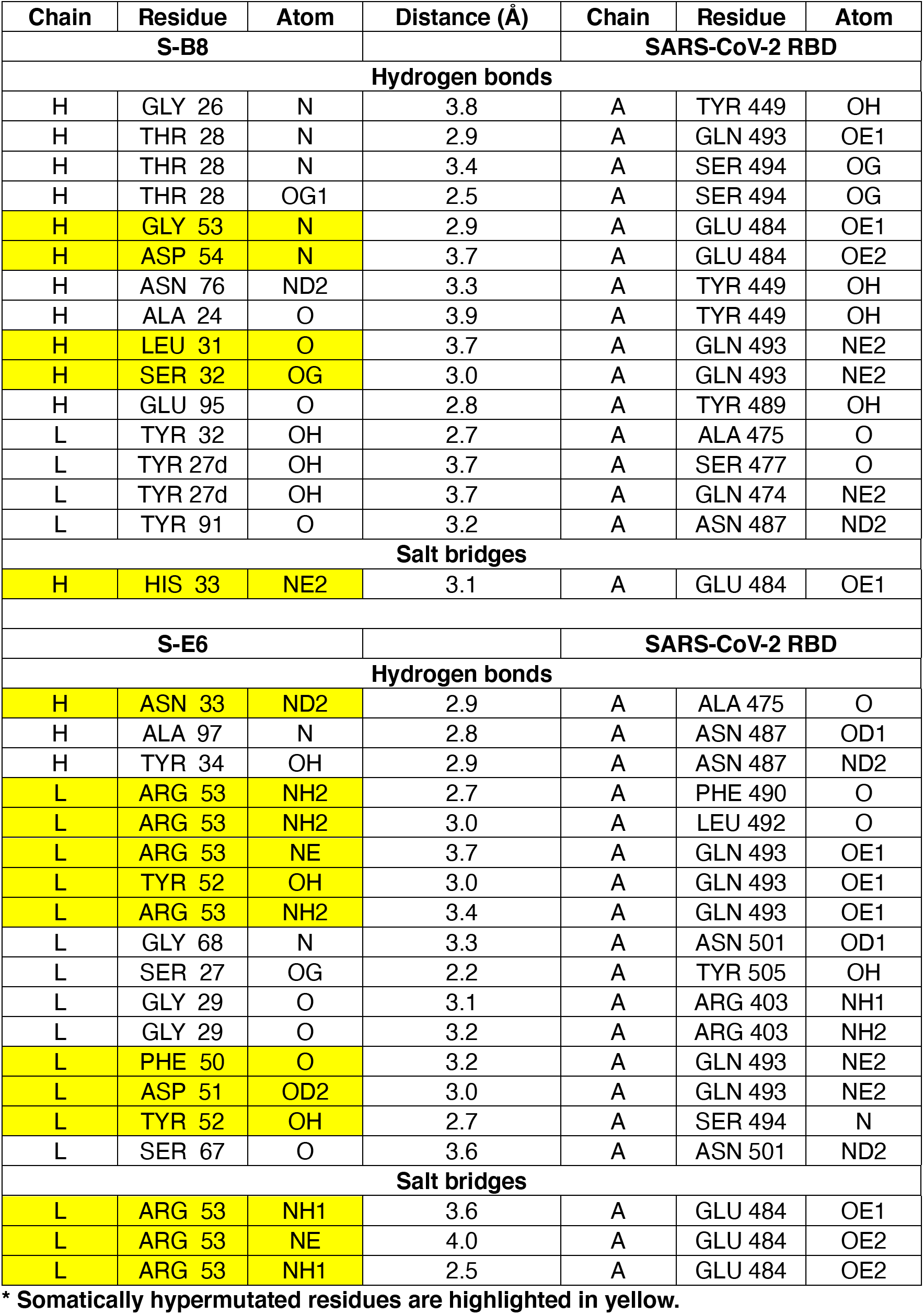
Hydrogen bonds and salt bridges identified at the antibody-RBD interface using the PISA program*.

## REFERENCES

1. N. Vabret et al., Immunology of COVID-19: current state of the science. Immunity 52, 910–941 (2020).

2. E. O. Saphire et al., Systematic analysis of monoclonal antibodies against Ebola virus GP defines features that contribute to protection. Cell 174, 938–952 (2018).

3. T. F. Rogers et al., Isolation of potent SARS-CoV-2 neutralizing antibodies and protection from disease in a small animal model. Science 369, 956–963 (2020).

4. A. Z. Wec et al., Broad neutralization of SARS-related viruses by human monoclonal antibodies. Science 369, 731–736 (2020).

5. J. Hansen et al., Studies in humanized mice and convalescent humans yield a SARS-CoV-2 antibody cocktail. Science 369, 1010–1014 (2020).

6. S. Ravichandran et al., Antibody signature induced by SARS-CoV-2 spike protein immunogens in rabbits. Sci Transl Med 12, eabc3539 (2020).

7. X. Chi et al., A neutralizing human antibody binds to the N-terminal domain of the Spike protein of SARS-CoV-2. Science 369, 650–655 (2020).

8. P. J. M. Brouwer et al., Potent neutralizing antibodies from COVID-19 patients define multiple targets of vulnerability. Science 369, 643–650 (2020).

9. D. F. Robbiani et al., Convergent antibody responses to SARS-CoV-2 in convalescent individuals. Nature 584, 437–442 (2020).

10. Y. Cao et al., Potent neutralizing antibodies against SARS-CoV-2 identified by high-throughput single-cell sequencing of convalescent patients’ B cells. Cell 182, 73–84 e16 (2020).

11. D. Pinto et al., Cross-neutralization of SARS-CoV-2 by a human monoclonal SARS-CoV antibody. Nature 583, 290–295 (2020).

12. C. Wang et al., A human monoclonal antibody blocking SARS-CoV-2 infection. Nat Commun 11, 2251 (2020).

13. L. Hanke et al., An alpaca nanobody neutralizes SARS-CoV-2 by blocking receptor interaction. Nat Commun 11, 4420 (2020).

14. Y. Wu et al., Identification of human single-domain antibodies against SARS-CoV-2. Cell Host Microbe 27, 891–898 e895 (2020).

15. X. Zeng et al., Blocking antibodies against SARS-CoV-2 RBD isolated from a phage display antibody library using a competitive biopanning strategy. bioRxiv, 2020.2004.2019.049643 (2020).

16. F. Bertoglio et al., SARS-CoV-2 neutralizing human recombinant antibodies selected from pre-pandemic healthy donors binding at RBD-ACE2 interface. bioRxiv, 2020.2006.2005.135921 (2020).

17. B. Felding-Habermann et al., Combinatorial antibody libraries from cancer patients yield ligand-mimetic Arg-Gly-Asp-containing immunoglobulins that inhibit breast cancer metastasis. Proc Natl Acad Sci U S A 101, 17210–17215 (2004).

18. R. A. Lerner, Combinatorial antibody libraries: new advances, new immunological insights. Nat Rev Immunol 16, 498–508 (2016).

19. J. Xie, H. Zhang, K. Yea, R. A. Lerner, Autocrine signaling based selection of combinatorial antibodies that transdifferentiate human stem cells. Proc Natl Acad Sci U S A 110, 8099–8104 (2013).

20. Z. Yang et al., A cell-cell interaction format for selection of high-affinity antibodies to membrane proteins. Proc Natl Acad Sci U S A 116, 14971–14978 (2019).

21. T. Zheng et al., Antibody selection using clonal cocultivation of *Escherichia coli* and eukaryotic cells in miniecosystems. Proc Natl Acad Sci U S A 115, E6145–E6151 (2018).

22. M. Qiang et al., Selection of an ASIC1a-blocking combinatorial antibody that protects cells from ischemic death. Proc Natl Acad Sci U S A 115, E7469–E7477 (2018).

23. C. S. Gao, O. Brummer, S. L. Mao, K. D. Janda, Selection of human metalloantibodies from a combinatorial phage single-chain antibody library. J Am Chem Soc 121, 6517–6518 (1999).

24. See MATERIALS AND METHODS section.

25. C. W. Tan et al., A SARS-CoV-2 surrogate virus neutralization test based on antibody-mediated blockage of ACE2-spike protein-protein interaction. Nat Biotechnol 38, 1073–1078 (2020).

26. H. Yao et al., Patient-derived mutations impact pathogenicity of SARS-CoV-2. medRxiv, 2020.2004.2014.20060160 (2020).

27. J. Lan et al., Structure of the SARS-CoV-2 spike receptor-binding domain bound to the ACE2 receptor. Nature 581, 215–220 (2020).

28. R. L. Stanfield, I. A. Wilson, Antibody structure. Microbiol Spectr 2, AID-0012-2013 (2014).

29. I. A. Wilson, R. L. Stanfield, Antibody-antigen interactions. Curr Opin Struc Biol 3, 113–118 (1993).

30. J. Ye, N. Ma, T. L. Madden, J. M. Ostell, IgBLAST: an immunoglobulin variable domain sequence analysis tool. Nucleic Acids Res 41, W34–40 (2013).

31. N. C. Wu et al., An alternative binding mode of IGHV3-53 antibodies to the SARS-CoV-2 receptor binding domain. Cell Rep 33, 108274 (2020).

32. M. Yuan et al., Structural basis of a shared antibody response to SARS-CoV-2. Science 369, 1119–1123 (2020).

33. C. O. Barnes et al., SARS-CoV-2 neutralizing antibody structures inform therapeutic strategies. Nature, (2020).

34. M. Yuan, H. Liu, N. C. Wu, I. A. Wilson, Recognition of the SARS-CoV-2 receptor binding domain by neutralizing antibodies. Biochem Biophys Res Commun, (2020).

35. R. A. Lerner, Rare antibodies from combinatorial libraries suggests an S.O.S. component of the human immunological repertoire. Mol Biosyst 7, 1004–1012 (2011).

36. C. Kreer et al., Longitudinal isolation of potent near-germline SARS-CoV-2-neutralizing antibodies from COVID-19 patients. Cell 182, 843–854 e812 (2020).

37. E. Seydoux et al., Analysis of a SARS-CoV-2-infected individual reveals development of potent neutralizing antibodies with limited somatic mutation. Immunity 53, 98–105 (2020).

38. Y. Weisblum et al., Escape from neutralizing antibodies by SARS-CoV-2 spike protein variants. eLife 9, e61302 (2020).

39. L. Xu et al., Design and characterization of a human monoclonal antibody that modulates mutant Connexin 26 hemichannels implicated in deafness and skin disorders. Front Mol Neurosci 10, 298 (2017).

40. S. Xia et al., Inhibition of SARS-CoV-2 (previously 2019-nCoV) infection by a highly potent pan-coronavirus fusion inhibitor targeting its spike protein that harbors a high capacity to mediate membrane fusion. Cell Res 30, 343–355 (2020).

41. K.-Y. A. Huang et al., Breadth and function of antibody response to acute SARS-CoV-2 infection in humans. bioRxiv, 2020.2008.2028.267526 (2020).

42. D. C. Ekiert et al., A highly conserved neutralizing epitope on group 2 influenza A viruses. Science 333, 843–850 (2011).

43. M. Yuan et al., A highly conserved cryptic epitope in the receptor binding domains of SARS-CoV-2 and SARS-CoV. Science 368, 630–633 (2020).

44. Z. Otwinowski, W. Minor, Processing of X-ray diffraction data collected in oscillation mode. Methods Enzymol 276, 307–326 (1997).

45. A. J. McCoy et al., Phaser crystallographic software. J Appl Crystallogr 40, 658–674 (2007).

46. H. Liu et al., Cross-neutralization of a SARS-CoV-2 antibody to a functionally conserved site is mediated by avidity. Immunity (2020), in press.

47. P. Emsley, K. Cowtan, Coot: model-building tools for molecular graphics. Acta Crystallogr D Biol Crystallogr 60, 2126–2132 (2004).

48. P. D. Adams et al., PHENIX: a comprehensive Python-based system for macromolecular structure solution. Acta Crystallogr D Biol Crystallogr 66, 213–221 (2010).

49. E. Krissinel, K. Henrick, Inference of macromolecular assemblies from crystalline state. J Mol Biol 372, 774–797 (2007).

